# Enhanced strand transfer and mismatch extension by HIV-1C reverse transcriptase promote sequence motif duplication

**DOI:** 10.64898/2026.03.05.709784

**Authors:** Arun Panchapakesan, Aditya Pradeepbhai Joshi, Afzal Amanullah, Neelam Pargain, Kavita Mehta, Manisenthil Shanmugam, Jayendra Singh, Chhavi Saini, Narendra Nala, Udupi A Ramagopal, Udaykumar Ranga

**Affiliations:** HIV-AIDS Laboratory, Molecular Biology and Genetics Unit, Jawaharlal Nehru Centre for Advanced Scientific Research, Jakkur, Bengaluru, Karnataka, India; Molecular Biology Laboratory, Y. R. Gaitonde Centre for AIDS Research and Education (YRGCARE), Chennai, Tamil Nadu, India; Flow Cytometry Facility, Molecular Biology and Genetics Unit, Jawaharlal Nehru Centre for Advanced Scientific Research, Jakkur, Bengaluru, Karnataka, India; Division of Biological Sciences, Poornaprajna Institute of Scientific Research, Poornaprajnapura, Bidalur, Bengaluru, India; Department of Microbiology and FST, School of Science, GITAM University, 530045, Visakhapatnam, India

**Keywords:** HIV-1C, Reverse transcriptase, Nonhomologous recombination, sequence duplication

## Abstract

Genetic diversification of HIV-1 is driven largely by the error-prone activity of reverse transcriptase (RT) and frequent template switching during reverse transcription. A rare outcome of nonhomologous recombination is sequence motif duplication, which can alter viral gene regulation and protein function. Previous studies have shown that such duplications occur at significantly higher frequencies in HIV-1 subtype C (HIV-1C), particularly within the long terminal repeat (LTR) and p6-Gag regions, where they can confer replication advantages. However, the mechanistic basis for this subtype-specific bias remains unclear. We therefore investigated whether intrinsic biochemical properties of HIV-1C RT contribute to its elevated duplication frequency. Bioinformatic analysis of 6,877 full-length HIV-1 genomes identified four duplication hotspots, with the highest frequencies in HIV-1C. Comparative sequence analysis of RT revealed several subtype-specific residues, including a highly conserved threonine at position 359 (T359) in the connection domain of HIV-1C RT. Structural modeling suggested that T359 can form an additional hydrogen bond with the nascent cDNA, potentially stabilizing the RT-template complex. Biochemical characterization of recombinant RT variants demonstrated that residue 359 modulates polymerase activity and maintains subtype-specific optimal catalytic function. Functional assays further revealed that HIV-1C RT exhibits enhanced template strand transfer compared with HIV-1B RT. In addition, next-generation sequencing-based primer extension assays showed that HIV-1C RT extends mismatched 3′ termini more efficiently across multiple mismatch types. Together, these findings indicate that subtype-specific biochemical properties of HIV-1C RT, particularly enhanced strand transfer and mismatch extension mediated in part by T359, promote nonhomologous recombination events that generate sequence motif duplications. This work provides a mechanistic explanation for the elevated duplication frequency characteristic of HIV-1C and highlights how subtle RT polymorphisms can shape viral evolutionary trajectories.

## Introduction

The extraordinary genetic diversity of Human Immunodeficiency Virus Type-1 (HIV-1), which underlies its capacity for rapid adaptation and persistence within the host, arises largely from the activity of the viral reverse transcriptase (RT). Unlike most cellular polymerases, HIV-1 RT lacks a 3′–5′ exonuclease proofreading function, resulting in frequent nucleotide misincorporation during DNA synthesis. However, its intrinsic error rate is comparable to that of other retroviral RTs (Hu and Hughes, 2012). In addition to mutations, the exceptionally high frequency of recombination, driven by frequent template switching during reverse transcription, plays a central role in generating HIV-1 genetic diversity (Onafuwa-Nuga & Telesnitsky, 2009).

Recombination, an essential feature of retroviral replication that drives viral evolution and adaptation, typically occurs between homologous regions of co-packaged RNA genomes during reverse transcription. A small proportion (0.1–1%) of recombination events, however, arise through nonhomologous pathways involving microhomology domains (Zhang & Temin, 1993). In such cases, the site at which RT dissociates from the donor RNA and the site at which it resumes synthesis on the acceptor RNA are not identical, resulting in sequence duplications, deletions, or insertion–deletion events (Onafuwa-Nuga & Telesnitsky, 2009).

Among these outcomes, sequence duplications are particularly intriguing because they can modulate viral gene expression, protein structure, and replication capacity. Although many duplication events are likely deleterious and eliminated by purifying selection, those that confer a fitness advantage can persist and expand within viral populations. Recurrent duplications have been reported in specific regions of the HIV-1 genome, most notably within the long terminal repeat (LTR) and the p6-Gag region, and are associated with enhanced replicative fitness (Bachu, Yalla, et al., 2012; Martins et al., 2015; Sharma et al., 2018). Duplication of the NF-κB motif in the LTR enhances transcriptional activity (Bachu, Mukthey, et al., 2012), whereas duplication of the PTAP motif in p6-Gag can compensate for deleterious immune-escape or drug-resistance mutations (Sharma et al., 2017). Strong selective pressures on these regions have therefore rendered them prominent duplication ‘hotspots’ in HIV-1 sequence databases.

Sequence duplications may also influence antiretroviral drug resistance. Several studies have documented higher frequencies of PTAP duplications among drug-resistant HIV-1 strains (Brindeiro et al., 2002; Martins et al., 2011; Peters et al., 2001; Tamiya et al., 2004). In HIV-1 subtype C (HIV-1C), PTAP duplication frequencies are higher in both drug-naïve (23%) and treatment-failure (54%) cohorts compared to subtypes B and F (Martins et al., 2011). PTAP duplication has been proposed to enhance proteolytic processing at the NC-Sp2-p6 junction, thereby compensating for fitness deficits imposed by resistance mutations (Martins et al., 2015). These observations suggest an interplay between subtype-specific viral determinants, sequence duplication, and the evolution of drug resistance.

Notably, motif duplications in both the LTR and gag regions occur more frequently in HIV-1C than in other subtypes. Among full-length viral genomes in extant databases, LTR duplications are observed in 27.3% of HIV-1B sequences and 34.5% of HIV-1C sequences. Similarly, PTAP duplications are present in 16.9% of HIV-1B and 30.4% of HIV-1C sequences (Bachu, Mukthey, et al., 2012; Sharma et al., 2017). These findings raise the possibility that HIV-1C has an intrinsic propensity to generate sequence motif duplications, potentially reflecting subtype-specific differences in viral replication. The molecular basis for this disparity, however, remains unclear.

Because recombination is a prerequisite for sequence duplication, RT function is a logical focus for mechanistic investigation. HIV-1 RT mediates template strand transfer during reverse transcription, and its recombination profile is influenced by intrinsic properties such as polymerase activity, processivity, fidelity, and RNase H activity (Hwang et al., 2001; Lanciault & Champoux, 2006; Onafuwa-Nuga & Telesnitsky, 2009). For example, nucleotide misincorporation can slow polymerization and promote recombination (Chin et al., 2007; Palaniappan et al., 1996; Schlub et al., 2014). Although biochemical analyses suggest broadly similar enzymatic parameters for HIV-1B and HIV-1C RTs (Xu et al., 2010), studies using chimeric viral backbones have revealed subtype-specific differences in replication kinetics (Iordanskiy et al., 2010). Thus, even in the context of conserved biochemical properties, HIV-1C RT may employ distinct molecular mechanisms that favor higher frequencies of recombination and duplication. Sequence duplications, though relatively rare, offer a selective snapshot of these processes, as they are retained when they confer a replication advantage.

The present study addresses this gap by examining how subtype-specific polymorphisms in HIV-1C RT, particularly the T359 residue, may contribute to elevated sequence motif duplication. Through integrated bioinformatic, biochemical, and molecular analyses, we aim to define the mechanistic basis of sequence duplication in HIV-1C and to clarify its implications for viral evolution, replicative fitness, and drug resistance.

## Results

### Four hotspots of sequence motif duplication exist in the HIV-1C genome

Our previous work identified hotspots of sequence motif duplication in two regions of the HIV-1C genome, the LTR and p6 (Bachu, Yalla, et al., 2012; Sharma et al., 2018). This observation prompted us to examine whether similar duplication hotspots occur elsewhere in the HIV-1C genome. We therefore retrieved 6,877 full-length HIV-1 genomes representing subtypes A1, B, C, D, F1, and G from the Los Alamos National Laboratory (LANL) HIV Sequence Database. Using a custom Python script, we systematically mapped sequence duplications across these genomes. In total, we identified 389 distinct loci with sequence duplications, distributed broadly across the genome, with an average inter-duplication distance of ∼24.9 bp. Two notable exceptions were p24, where no duplications were detected, and pol, where only two events were observed.

The apparent scarcity of duplications in p24 and pol is likely influenced by sampling bias in database-derived datasets. Because duplications frequently disrupt reading frames or impair protein function, such variants would be subject to strong negative selection in vivo and are therefore unlikely to be amplified or sequenced using standard approaches. Conversely, duplications that confer a selective advantage, such as those in the LTR and p6 regions, are more likely to persist and thus appear overrepresented. Consistent with this expectation, we observed a marked enrichment of duplications in these regions, forming clear hotspots. Despite this limitation, the analysis indicates that duplications can arise throughout the HIV-1 genome but are preferentially concentrated at specific loci, likely reflecting selective advantages. In addition to the previously reported hotspots in the LTR and p6 regions, we identified two additional hotspots in the envelope and nef regions, where duplications were consistently observed across all subtypes (Figures 1B and 1C).

**Figure 1.**
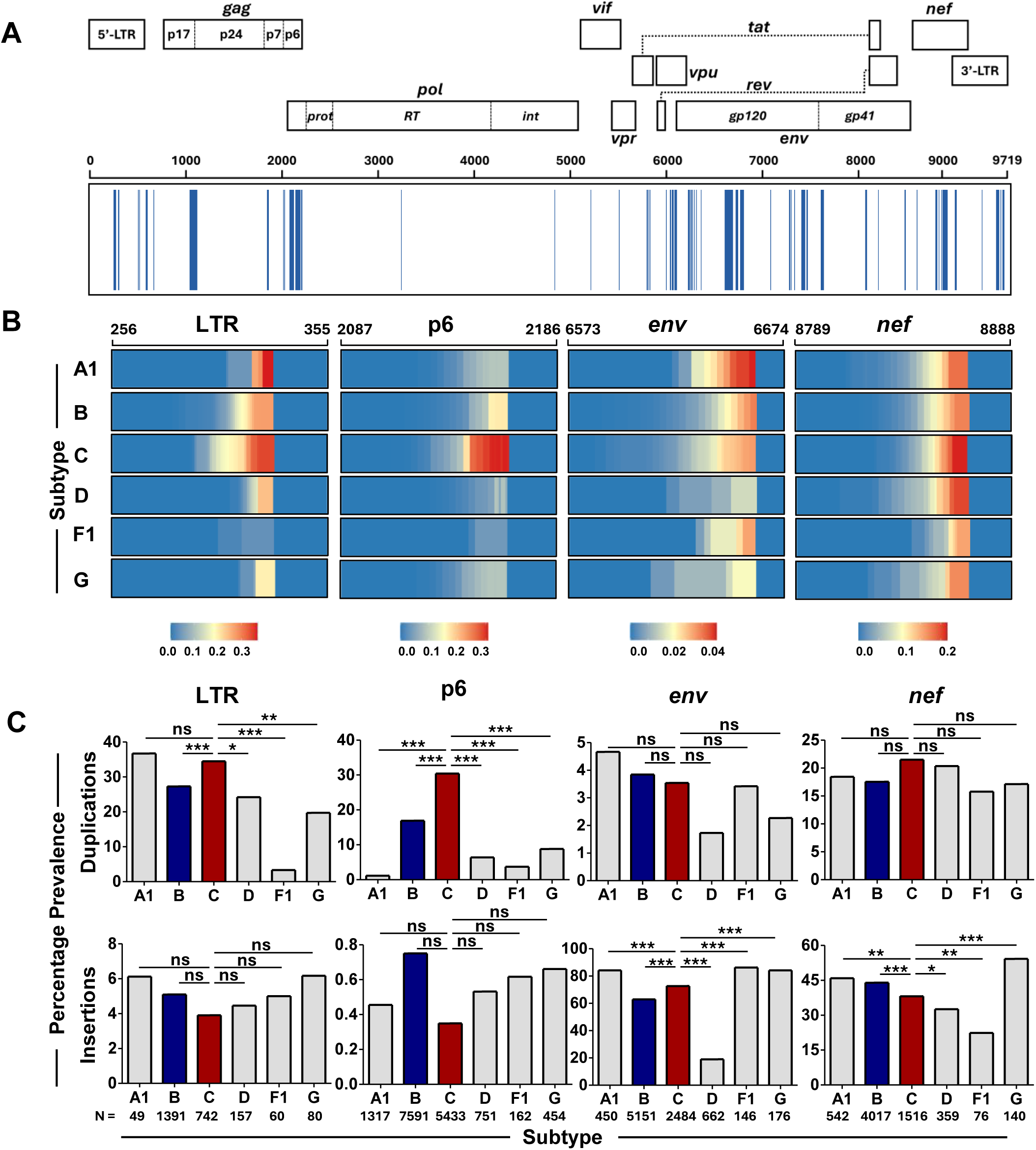
HIV-1C duplicates sequences at a higher frequency at hotspots. (A) Sequence duplications are observed throughout the HIV-1 genome. Full-length HIV-1 genome sequences (n=6,877) representing all major subtypes were downloaded from the HIV-LANL database and analyzed using a custom Python script to identify duplication events. A duplication at a given location is shown using vertical blue lines. Genomic positions are indicated using HXB2 coordinates. (B) HIV-1C exhibits a high frequency of duplications across known hotspots. Sequences of the LTR, p6, env, and nef regions from subtypes A1, B, C, D, F1, and G were downloaded from the HIV-LANL database and screened for sequence duplications. The heatmap illustrates the per-base frequency of duplications across a 100 base pair region in each location, as depicted using HXB2 coordinates. (C) HIV-1C shows significantly higher duplication frequencies in the LTR and p6. Bar graphs showing the proportion of sequences in the database that contain duplications (top) and insertions (bottom) in the four hotspots in different subtypes. Subtypes B and C are highlighted. Statistical analysis was performed using Pearson’s Chi-square test. * p < 0.05, ** p < 0.01, *** p < 0.001.. * p < 0.05, ** p < 0.01, *** p < 0.001. N represents the number of sequences used in the analysis.

Among the subtypes analyzed, HIV-1C showed the highest duplication frequencies in the LTR, p6, and nef regions, accounting for 34.5%, 30.5%, and 21.5% of duplications, respectively. The envelope region was an exception: subtypes A1 and B showed slightly higher duplication frequencies (4.6% and 3.8%) than HIV-1C (3.5%) (Figures 1B and 1C). Overall, the average per-base duplication frequency across the four hotspots was markedly higher in HIV-1C than in other subtypes (Figure 1B). However, only the LTR and p6 hotspots in HIV-1C reached statistical significance (Figure 1C). One possible explanation is that insertions in these two regions represent true motif duplications that confer a replication advantage, as previously demonstrated (Bachu, Yalla, et al., 2012; Martins et al., 2015; Sharma et al., 2018).

In both regions, sequence duplications represented the predominant class of insertions across all subtypes. On average, duplications accounted for 24.3% and 11.2% of sequences in the LTR and p6 regions, respectively, whereas other insertions were comparatively rare (5.1% and 0.5%) (Figure 1C, bottom panel). In contrast, the envelope and nef regions showed the opposite pattern: although insertions were present, most did not represent duplication of adjacent sequence motifs (Figure 1C, bottom panel). This suggests that insertions in these regions arise largely from stochastic variation or structural tolerance rather than motif-specific duplication. Collectively, these analyses indicate that sequence motif duplication occurs more frequently in HIV-1C, with duplications in the LTR and p6 regions conferring a replication fitness advantage.

### The T359 signature amino acid residue can form an additional hydrogen bond with the cDNA

To determine whether the higher frequency of sequence duplications in HIV-1C reflects subtype-specific features of reverse transcriptase (RT), we performed a comparative analysis of HIV-1 RT sequences downloaded from the LANL database. One full-length genome per drug-naïve individual was included, and sequences were grouped based on copy-number variation at three loci: the NF-κB (3 vs. 4 copies) and RBEIII (1 vs. 2 copies) elements in the LTR, and the PTAP motif (1 vs. 2 copies) in p6-Gag.

Using this dataset, we performed Viral Epidemiology Signature Pattern Analysis (VESPA) on 100 randomly selected RT sequences representing each of the major HIV-1 subtypes A1, B, C, and D. Subtypes F1 and G were excluded because of insufficient numbers of p66 sequences in the database. This analysis identified 28 amino acid residues specific to HIV-1C (Table 1). Applying a frequency threshold of >75% in HIV-1C and <25% in other subtypes reduced this set to six candidate residues: A36, T48, A200, Q245, T359, and R530. These residues span the fingers (A36, T48), palm (A200), thumb (Q245), connection (T359), and RNase H (R530) domains of RT (Table 1).

**Table 1:**
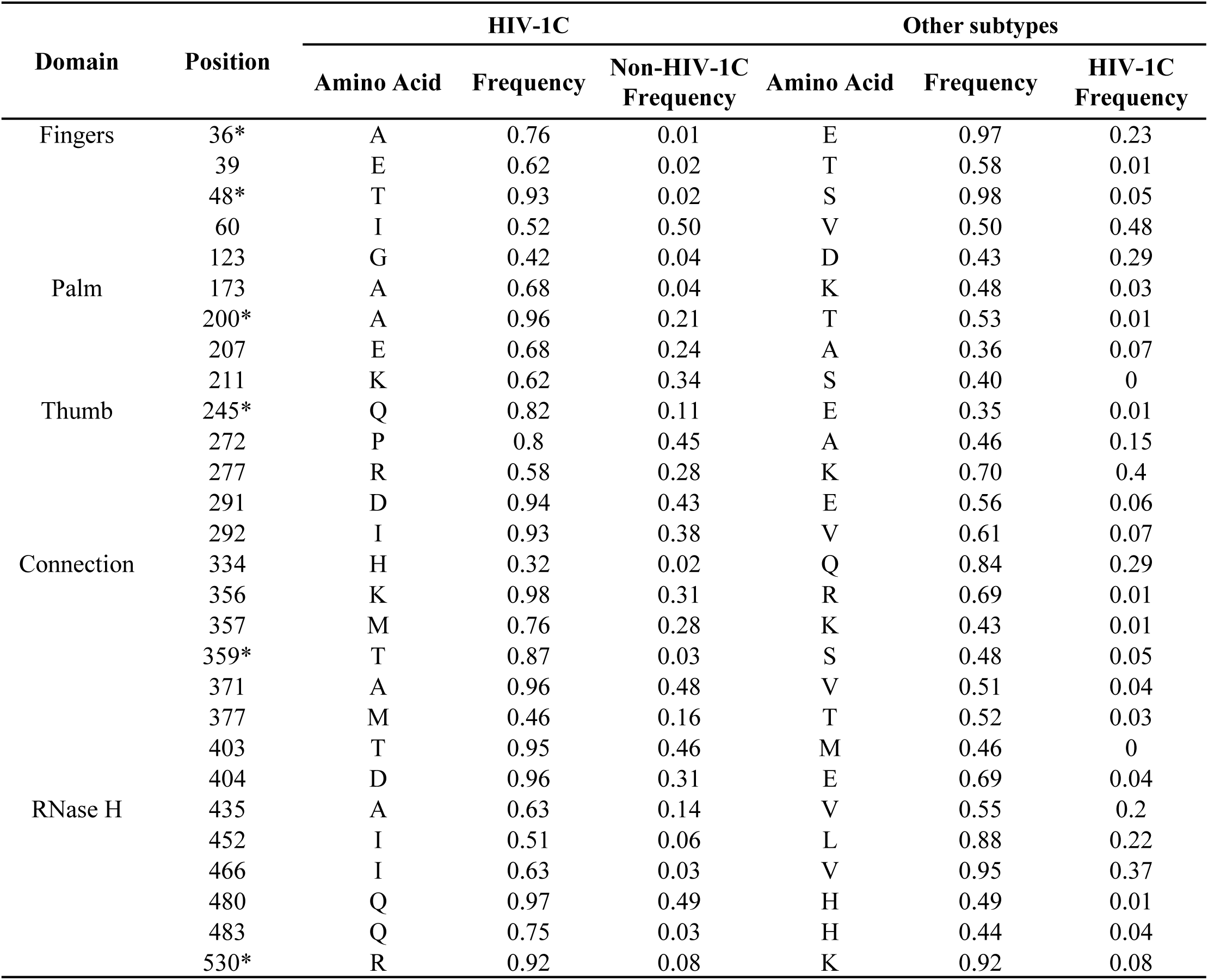
The frequencies of the Signature Amino acid residues in HIV-1C Reverse Transcriptase. HIV-1C RT sequences were subjected to signature amino acid residue analysis using the VESPA tool available at the HIV-LANL database with subtypes A1, B, and D as background. Subtypes F1 and G were not included in the analysis due to the low number of sequences available. The single letter amino acid code is used. Amino acid residues selected for subsequent analyses are shown with an Asterix.

Among these positions, T359 was of particular interest for three reasons. First, its presence in HIV-1C is exclusive to this subtype, as is that of two other residues at positions 36 and 48. In contrast, the three corresponding residues at the other three positions: 200, 245, and 530, occurred at comparatively lower frequencies in other subtypes (Figure 2A, Supplementary Table S1). Second, the glycine-to-threonine substitution at position 359 is non-conservative, introducing increased polarity and the potential for hydrogen bond formation. Four residues, alanine, glycine, serine, and threonine, occur at this position across subtypes with distinct distributions (Figure 2A, Supplementary Table S1). Glycine predominates in subtypes B and D (92.3% and 91.2%), serine in A1 (88.8%), whereas threonine is the dominant residue in HIV-1C (83.7%) and is nearly absent from other subtypes. Third, structural mapping of the six residues onto the RT structure showed that T359 lies near the nucleic acid-binding cleft, suggesting potential interaction with the DNA/RNA substrate (Figure 2B).

**Figure 2:**
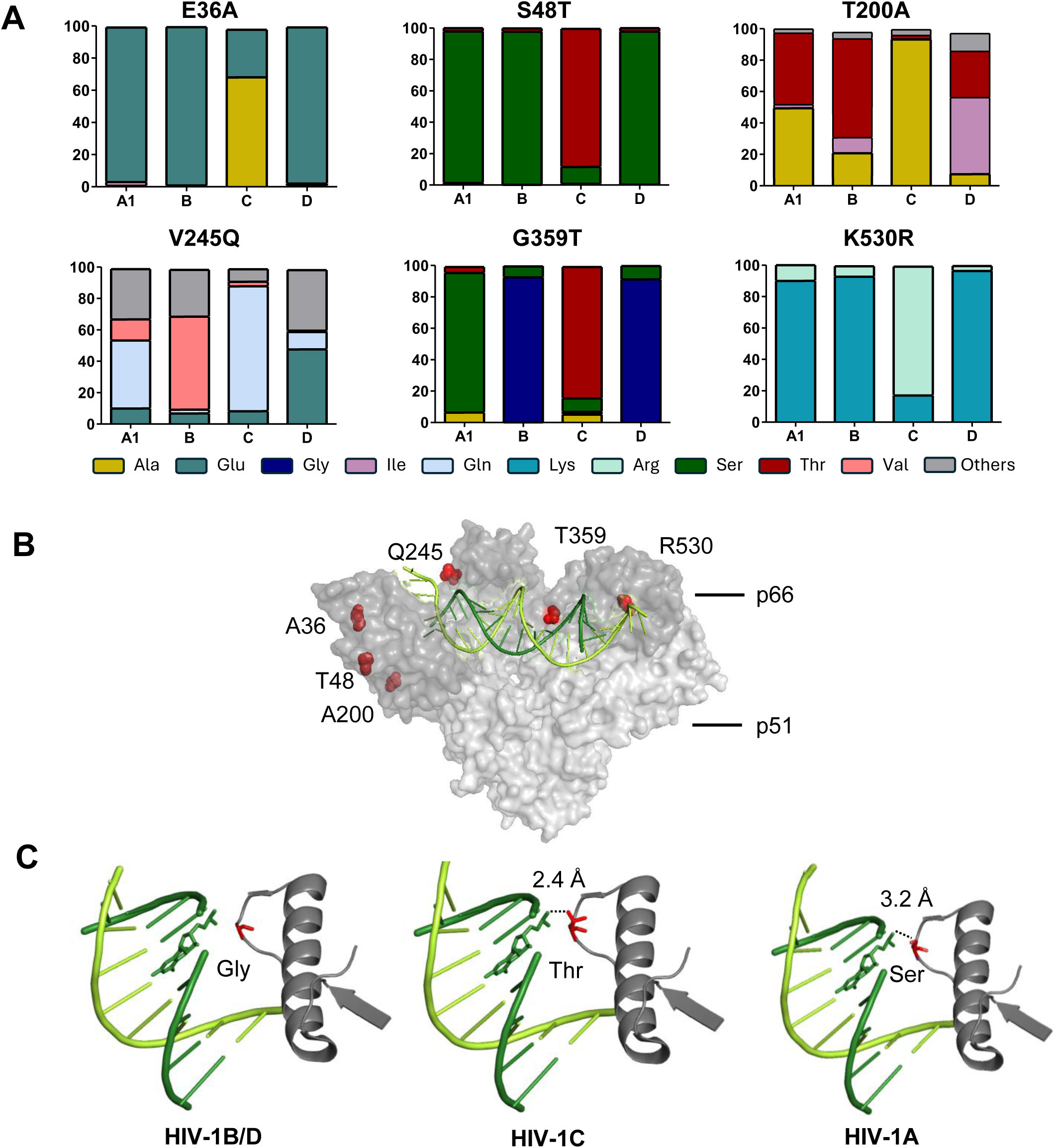
The T359 Residue of HIV-1C forms a hydrogen bond with the cDNA. (A) Amino acid variation at six signature residues in HIV-1C reverse transcriptase. Amino acid frequencies at six key positions identified through sequence analysis. The reference amino acids indicated in the panel headings correspond to HXB2 and Indie-C1, respectively. **(B) Spatial proximity of the T359 residue to the DNA strand in HIV-1 RT.** A high-resolution structure of HIV-1B RT (PDB ID: 5J2M) was visualized using PyMOL. The six residues of interest are depicted in red as enlarged spheres. The DNA template and cDNA strands are shown in light and dark green, respectively. The p66 and p51 subunits are coloured dark and light grey, respectively and are shown in the background. **(C) Hydrogen bond formation between the T359 residue and cDNA.** The structural environment around residue 359 was examined using the same RT structure. The native Glycine (left), and modified Threonine (centre) and Serine substitutions (right) at this position were modelled. The cDNA base involved in hydrogen bonding is shown as sticks. The distance between the hydroxyl group of the residue and the phosphate group of the cDNA was measured in PyMOL and is represented by a dotted line.

To evaluate this possibility, we modeled serine and threonine substitutions using the HIV-1B RT crystal structure (PDB 5J2M) as a template. In subtypes B and D, which contain glycine at position 359, the absence of a side chain prevents strong interactions with DNA. In contrast, in subtypes A1 and C, which harbour serine and threonine, respectively, the hydroxyl groups of these residues lie close to a phosphate atom on the nascent cDNA backbone, enabling potential hydrogen-bond formation. The predicted atomic distances are ∼3.2 Å for serine and ∼2.4 Å for threonine (Figure 2C). The shorter distance for threonine suggests a stronger interaction and supports a possible role for this residue in modulating RT-template contacts in HIV-1C.

### The T359 residue ensures an optimal RT activity in HIV-1C

Sequence and structural analyses indicated that the threonine residue at position 359 of HIV-1C RT is highly conserved and can form an additional hydrogen bond with the nascent DNA, suggesting a potential effect on polymerase function. Because polymerase activity influences downstream processes such as strand transfer, recombination, and sequence duplication, we examined how variation at residue 359 affects diverse RT functions.

To address this question, we constructed two panels of RT variants using HIV-1B and HIV-1C RTs as backbones. Each panel contained three enzymes differing only at residue 359, which carried the naturally occurring amino acids glycine, serine, or threonine (Figure 2A). Recombinant RTs were generated by expressing the p66 and p51 subunits separately in *E. coli* M15 cells. The proteins were purified by sequential Ni-NTA and ion-exchange chromatography and assembled in vitro for biochemical analyses.

Polymerase activity was measured by ^32^P-dATP incorporation on a poly-U template over a kinetic time course. All six enzymes were active and showed progressive incorporation of the isotope (Figure 3A). However, clear subtype-dependent differences in polymerase kinetics emerged. HIV-1B RTs (B-RTs) containing glycine or serine, the most common residues at this position in this subtype, displayed substantially higher activity than the threonine-containing variant. At 30 min, glycine- and serine-containing B-RTs incorporated 27.47 ± 1.11 and 26.26 ± 2.85 pmol dATP, respectively, whereas the threonine variant incorporated only 10.81 ± 1.16 pmol. In contrast, HIV-1C RT (C-RT) containing the canonical T359 showed the lowest catalytic activity, which increased markedly upon substitution. After 30 min, activity rose from 17.1 ± 1.79 pmol in the T359 enzyme to 179.28 ± 7.66 pmol and 54.18 ± 5.19 pmol with glycine and serine substitutions, respectively (Figure 3A). Thus, RTs carrying the canonical residue at position 359 for their subtype exhibited comparable activity; substitution with a non-canonical residue reduced activity in HIV-1B (threonine) but markedly increased it in HIV-1C (glycine).

**Figure 3.**
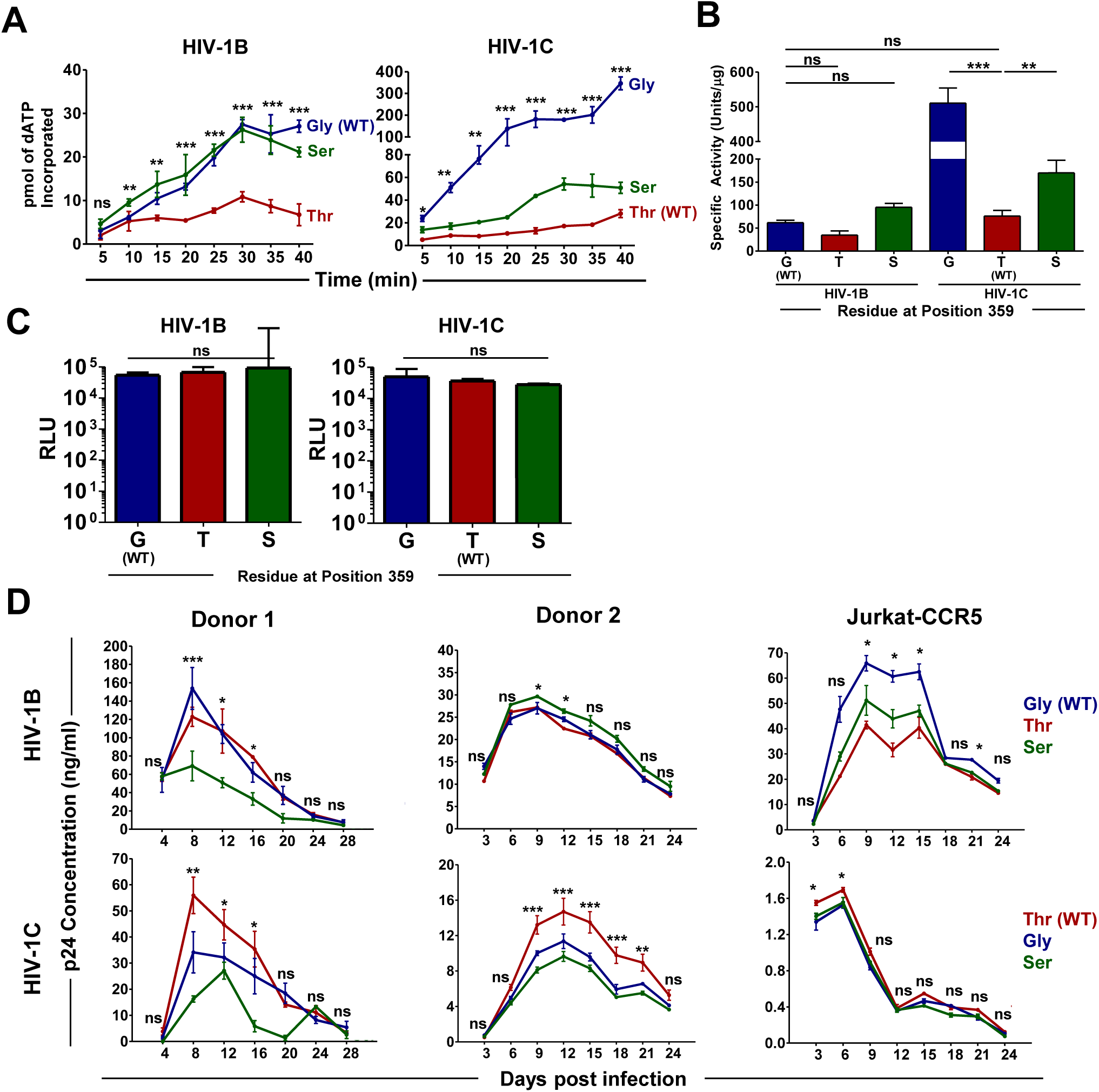
Polymerase function of the RT Variants. (A) cDNA synthesis rate of RT variants. cDNA synthesis rates of HIV-1B (left) and HIV-1C (right) RT variants expressed and purified from *E. coli.* A homopolymeric template, polyuridylic acid, was extended with a primer and ³²P-dATP incorporation was monitored by liquid scintillation at 5-min intervals. Data are presented as Mean ± SD and are representative of three independent experiments. Note the difference in the y-axis between subtypes. Data was analysed by two-way ANOVA with Bonferroni post hoc test. **(B) Specific activities of the RT variant proteins.** The amount of radioactive dATP incorporated in 10 min was used to calculate the specific activity of all RT variants. The data are representative of three independent experiments, and are plotted as Mean ± SD, followed by one-way analysis of variance and Tukey-Kramer post. hoc tests. **(C) Single-round infectivity of RT variant viral strains.** Replication-competent NL4-3 (HIV-1B) and Indie (HIV-1C) clones carrying Gly, Ser, or Thr at RT position 359 were used for the assay. Viral stock titers were measured by infecting TZM-bl cells with 10 ng p24 of each variant. Luciferase induction by Tat-driven LTR activity served as a surrogate for viral replication. Data represent Mean ± SD (n=3), analyzed by one-way ANOVA with Tukey–Kramer post hoc test. **(D) Replication profiles of RT variant viral strains.** Replication kinetics of RT-variant viral strains in Jurkat-CCR5 T-cells and CD8-depleted PBMCs from two donors. 3 x 10^6^ cells were infected with 10 ng p24 equivalent of each of the viral stocks and viral proliferation was estimated by measuring the p24 production at 3- or 4-day intervals using a commercial kit. The amino acid colour code is consistent across the figure. The data is represented as Mean ± SD was analysed by two-way ANOVA with Bonferroni post hoc test. ***p < 0.001, **p < 0.01, *p < 0.05, ns = not significant.

Subtype-specific differences were also evident in specific activity across the RT panel, consistent with previous reports (Iordanskiy et al., 2010; Xu et al., 2010). HIV-1B and HIV-1C RTs containing their canonical residues (G359 and T359, respectively) showed comparable activities of 64.98 ± 5.33 and 76.15 ± 12.24 U/μg (Figure 3B). In B-RT, substitution with serine moderately increased activity (95.24 ± 8.23 U/μg), whereas threonine reduced it to 34.65 ± 9.64 U/μg. In contrast, substitutions in C-RT produced striking increases: the canonical T359 enzyme (76.15 ± 12.24 U/μg) rose to 510.66 ± 43.39 U/μg (6.7-fold) and 170.25 ± 27.23 U/μg (2.23-fold) with glycine or serine substitutions, respectively (Figure 3B). These findings indicate that variation at position 359 strongly influences C-RT function but has only modest effects on B-RT.

The increase in catalytic activity observed upon substituting threonine with glycine or serine in C-RT suggests that T359 normally imposes a regulatory constraint on HIV-1C RT. Formation of an additional hydrogen bond between T359 and the nascent cDNA may slow catalytic kinetics, maintaining activity within an optimal range. Consistent with this model, replacing threonine with serine, which forms a weaker hydrogen bond due to a longer bond length (∼3.2 Å), produced only a modest increase in activity. In HIV-1, reverse transcription typically requires 8-33 hours depending on the host cell type (Murray et al., 2011); excessively rapid DNA synthesis could therefore compromise replication fitness. Notably, RT sequences vary substantially among HIV-1 subtypes. For example, the RTs of NL4-3 (HIV-1B) and Indie (HIV-1C), used in our assays, differ by 7.68% at the amino-acid level. Such intrinsic variation may confer higher catalytic potential to C-RT, which T359 may help restrain to prevent excessively rapid reverse transcription.

Because the T359G variant of HIV-1C RT showed the highest catalytic activity among the enzymes tested, we next evaluated the effects of residue 359 substitutions in replication-competent viruses. Viral panels analogous to the recombinant RT panels (Figure 3A) were generated using the NL4-3 (HIV-1B) and Indie-C1 (HIV-1C) molecular clones. Viral stocks produced in HEK293T cells showed comparable infectious titres across variants, as measured by TZM-bl luciferase assays following infection with 10 ng p24 (Figure 3C). Replication kinetics were then assessed in primary PBMCs from two healthy donors and in Jurkat-CCR5 cells. All variants replicated efficiently, with peak p24 production between days 6 and 12 irrespective of the backbone (Figure 3D). As expected, NL4-3-based viruses generally produced higher p24 levels than Indie-C1 derivatives.

Within each panel, the virus carrying the subtype-specific canonical residue at position 359 replicated most efficiently (Figure 3D). In the NL4-3 panel, the G359 strain (wild-type HIV-1B) reached 153.90 ± 39.63 ng/ml p24 at day 8 in donor 1 PBMCs, whereas the T359 and S359 variants produced 122.88 ± 18.15 and 69.09 ± 28.03 ng/ml, respectively. Similarly, in the Indie-C1 panel, the wild-type T359 virus produced 55.95 ± 12.09 ng/ml p24 at day 8, compared with 34.13 ± 13.64 and 16.25 ± 2.07 ng/ml for the G359 and S359 variants. Replication patterns were broadly consistent in donor 2 PBMCs and Jurkat-CCR5 cells, although modest donor-specific differences were observed, particularly in the NL4-3 panel. Thus, although the variant HIV-1C RTs containing non-canonical amino acid residues at position 359 showed higher catalytic activity than the enzyme containing the canonical threonine residue, in the context of the infectious viral clone, the viral strain containing the canonical threonine at position 359 demonstrated the highest level of proliferation, consistent with our model.

Together, these results demonstrate that residue 359 plays a key role in determining RT catalytic function. Canonical residues at this position, glycine in HIV-1B and threonine in HIV-1C, maintain optimal RT activity and are not interchangeable between subtypes. Notably, these residues occur at the highest frequencies within their respective subtypes and are rarely shared between them. Combined with the biochemical data, these findings suggest that efficient viral replication requires RT activity to remain within a defined functional window. Our results further indicate that T359 represents a subtype-specific regulatory adaptation in HIV-1C, moderating RT kinetics to maintain optimal activity and preserve replication fitness.

### HIV-1C RT mediates enhanced strand transfer in vitro

Because recombination is a prerequisite for sequence duplication, a higher recombination frequency could explain the greater prevalence of motif duplications in HIV-1C. Given the central role of residue T359 in HIV-1C RT and the likelihood that this residue forms an additional hydrogen bond with the nascent cDNA, we hypothesized that threonine may stabilize the RT-cDNA complex and thereby enhance recombination efficiency. However, previous experimental work using an HIV-1C molecular clone (Chin et al., 2005) reported findings inconsistent with this hypothesis. Importantly, that study used a subgenomic viral backbone, highlighting the need to reassess the model using a near-full-length viral construct.

To evaluate the effect of residue 359 on recombination, we employed a fluorescent EGFP reporter recombination assay (Levy et al., 2004; Rhodes et al., 2005). This assay measures template switching by quantifying cells expressing functional EGFP restored from two complementary frame-shifted defective EGFP precursors. Two viral panels were generated in the NL4-3 (HIV-1B) and Indie-C1 (HIV-1C) backbones, each containing glycine, serine, or threonine at position 359 (Figure 4A). In total, 18 pseudotypable, replication-competent viral constructs were produced (Figure 4A, Supplementary Table 2). Because the assay uses a single-round infection system, only one cycle of infection occurs, preventing additional rounds of recombination. We first confirmed that only constructs carrying an intact EGFP cassette, but not those containing premature termination codons, produced detectable fluorescence after transfection into HEK293 cells (Figure 4B).

**Figure 4.**
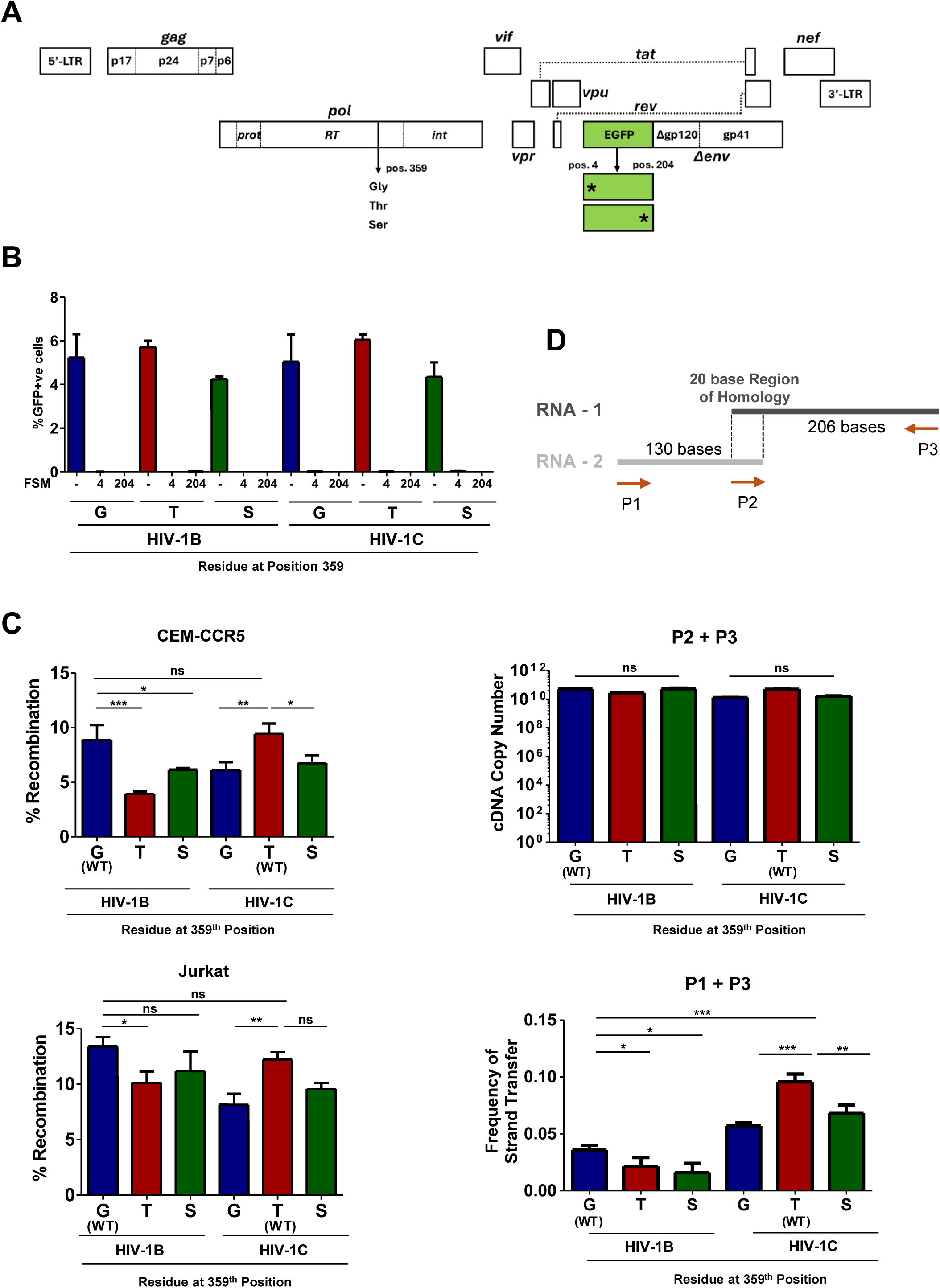
HIV-1B and -1C RTs recombine at comparable frequencies. (A) Schematic of the EGFP reporter virus design. Panels of three RT variants (Glycine, Threonine, or Serine at position 359) were generated using NL4-3 or Indie near full-length molecular clones. The viral envelope was replaced with an EGFP ORF, allowing single-round infection when pseudotyped. The EGFP reading frame contains frame-shift mutations (FSM) at amino acid positions 4 or 204, resulting in a total of 18 viral strains. **(B) Homozygous GFP mutant viruses do not express GFP.** Viral stocks of all 18 strains were produced in HEK293T cells and used to infect CEM-CCR5 or Jurkat cells. After 24 h, cells were activated with a cocktail of TNF-α (10 ng/ml), HMBA (10 ng/ml), and PMA (5 mM). GFP expression was assessed 24 h post-activation. The X-axis shows the position of the frameshift mutation. WT = Wild-type EGFP. **(C) Recombination frequencies of wild-type RTs.** Viral stocks with FSM4 and FSM204 EGFP mutations were used in different combinations to infect CEM-CCR5 or Jurkat cells, as shown in Supplementary Table 2. GFP expression was quantified as the percentage of GFP^+^ cells among infected cells. **(D) Strand-transfer assay schematic.** Two in vitro-transcribed RNA templates with a 20 bp overlap (dark/light grey lines) and primers (orange arrows) were used. Ct values were obtained via SYBR Green-based qPCR. **(E) HIV-1C RT variants show enhanced strand transfer.** cDNA copy numbers (primers P2+P3) were determined by regression (left) and used to normalize values of strand transfer PCR (P1+P3, right). Data are shown as mean ± SD from three independent experiments. Statistical significance was assessed using one-way ANOVA with Tukey–Kramer post hoc test. ***p < 0.001, **p < 0.01, *p < 0.05, ns = not significant.

Following co-infection with viruses encoding the two defective EGFP templates, the proportion of GFP-positive cells reflected the recombination capacity of each RT variant. Wild-type HIV-1B and HIV-1C RTs exhibited similar recombination frequencies in CEM-CCR5 cells, 8.86 ± 1.36% and 9.42 ± 0.94%, respectively (Figure 4C, top panel), consistent with earlier reports (Chin et al., 2005). Substitution of the canonical residue in either subtype significantly reduced recombination efficiency. In HIV-1B, replacing glycine at position 359 with threonine or serine reduced GFP-positive cells to 3.91 ± 0.21% and 6.14 ± 0.15%, respectively. Similarly, in HIV-1C, replacing the canonical threonine with glycine or serine yielded 6.09 ± 0.73% and 6.72 ± 0.73% GFP-positive cells.

A similar pattern was observed in Jurkat T cells, although overall recombination frequencies were slightly higher. For example, the GFP-positive fraction for the G359 HIV-1B RT decreased from 13.38 ± 0.85% to 10.1 ± 1.0% and 11.16 ± 1.77% after substitution with threonine or serine (Figure 4C, bottom panel). Likewise, replacing the canonical threonine in HIV-1C RT reduced GFP-positive cells from 12.02 ± 0.69% to 8.13 ± 0.99% and 9.56 ± 0.53%, respectively. These findings are consistent with the dynamic copy-choice model, in which faster reverse transcription correlates with reduced template switching.

Since sequence duplication requires RT to switch between the two genomic RNA templates, we next asked whether HIV-1C RT intrinsically promotes template switching more efficiently. To test this, we performed a template-switching assay using two single-stranded RNA templates with a 20-base overlap (Figure 4D, top panel). Successful strand transfer during reverse transcription produces a longer 336 bp cDNA product detectable by PCR, whereas the absence of strand transfer yields a shorter 206 bp fragment. The assay was performed using both HIV-1B and HIV-1C RT panels (Figure 4E). All six RT variants generated comparable levels of the 206 bp product that does not require template switching (Figure 4E, middle panel). However, marked differences, both between and within subtypes, were observed in the formation of the 336 bp product requiring strand transfer (Figure 4E, bottom panel).

Notably, all HIV-1C RT variants collectively exhibited higher strand-transfer activity than HIV-1B RTs. Within HIV-1C, the canonical T359 enzyme showed the highest switching frequency (0.096 ± 0.01), compared with the glycine and serine variants: 0.057 ± 0.00 and 0.07 ± 0.01, respectively (Figure 4E, bottom panel). In contrast, HIV-1B RT variants displayed substantially lower activity, with glycine (wild-type), threonine, and serine variants showing switching frequencies of 0.03 ± 0.00, 0.03 ± 0.00, and 0.02 ± 0.01, respectively. These results indicate that HIV-1C RT possesses intrinsically higher strand-transfer capacity, a key mechanistic driver of sequence duplication. Collectively, the data support a model in which T359 enhances template engagement through an additional hydrogen bond, whereas substitutions that weaken this interaction impair recombination-related functions. Thus, threonine at position 359 appears to represent a subtype-specific regulatory adaptation that finetunes reverse transcription dynamics to support the elevated duplication frequencies characteristic of HIV-1C.

### HIV-1C RT extends 3’-OH mismatches more efficiently

The phenomenon of sequence motif duplication requires that RT switch to the acceptor RNA template at a location that has already been copied from the donor RNA. However, such an erroneous strand switch, although very rare, imposes a constraint when it occurs, because the sequences at the growing end of the nascent cDNA may be discordant, thereby hampering efficient resumption of polymerization. Notably, retroviral RTs can anneal to mismatched sites using very short regions of complementarity, sometimes as few as three bases or fewer, termed micro-homology domains (MHDs), to continue reverse transcription. Such MHD-mediated annealing is predicted to be highly transient and unstable, making nonhomologous recombination events exceedingly rare (Zhang and Temin, 1993). Furthermore, following MHD-mediated hybridization, a base-pair mismatch at the 3′ end of the nascent cDNA is expected to occur in three out of four instances, requiring RT to efficiently extend a mispaired terminus to resume polymerization and complete the duplication event. For example, duplication of the NF-κB motif in HIV-1C results from extension of a mismatched C-A base pair (Figures 5A and B).

**Figure 5.**
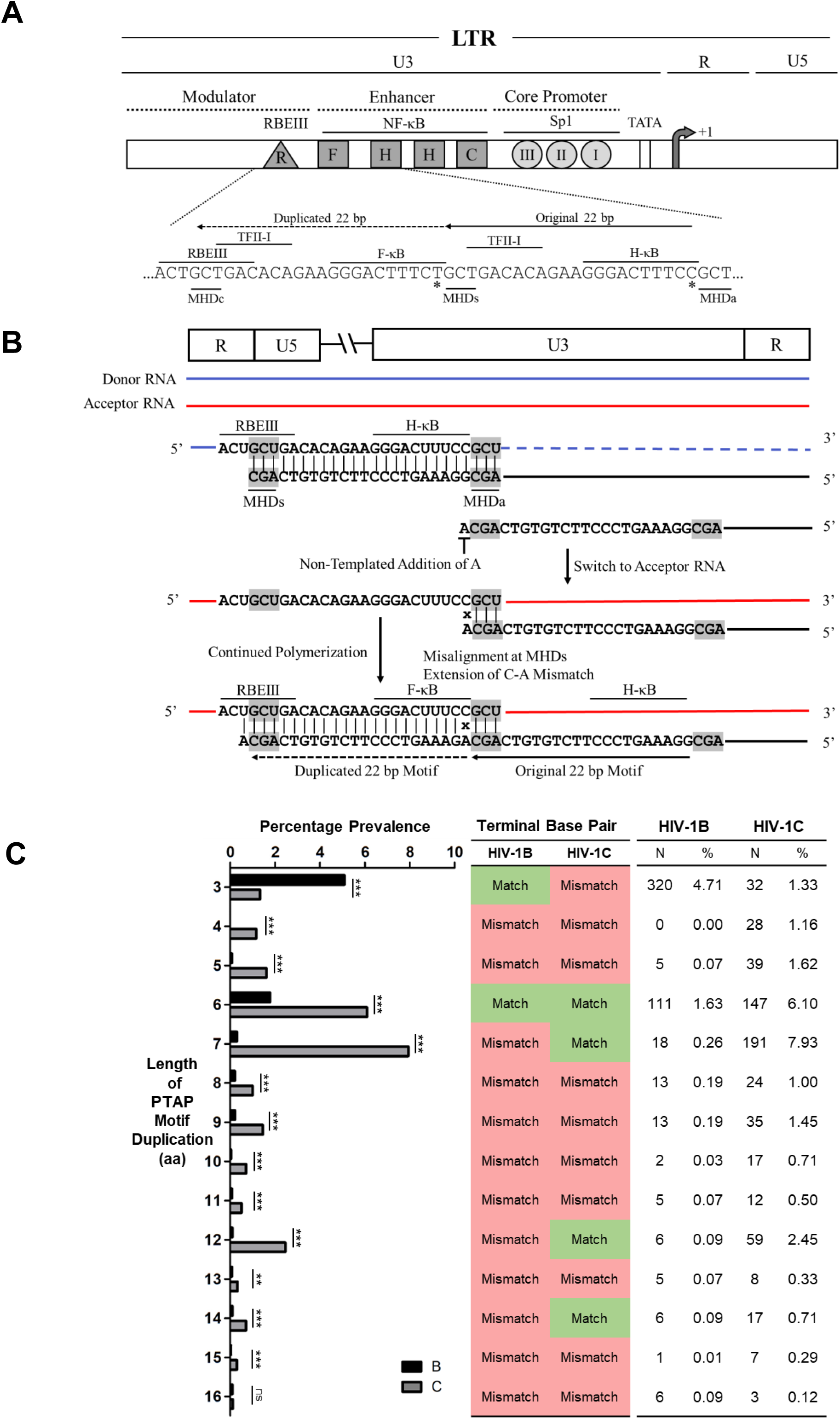
(A) Schematic representation of the 22 bp NF-κB motif duplication in HIV-1C mediated by extension of a mismatched base. (A) Organization of the HIV-1C LTR showing the relative positions of the four NF-κB motifs and other key transcription factor binding sites (TFBSs), along with their nucleotide sequences. Solid arrows indicate the original sequence, whereas dashed arrows denote the duplicated segment. The 3-base pair repeats flanking the original sequence, which is recreated in the duplication, called the Micro-homology domain (MHD), are underlined. The three such MHD triplets corresponding to positions where the RT stalls, re-aligns and creates a new copy are designated MHDs, MHDa, and MHDc, respectively. Asterisks mark the C-to-T variation distinguishing the H-κB and F-κB motifs. The duplicated 22 bp segment corresponds to positions 325–352 of the HXB2 reference sequence. Image reproduced from Panchapakesan and Ranga, Viruses, 2025 under the Creative Commons CC BY 4.0 license **(B) Schematic showing the template-switching mechanism and mismatch extension.** Donor and acceptor RNA templates are depicted in blue and red, respectively, while the nascent cDNA is shown in black. Dashed lines represent RNA regions degraded by the RNase H activity of reverse transcriptase (RT). The extended mismatched base is indicated by an “x” MHD motifs are shaded in gray. **(C) Spikes in sequence motif duplication are associated with perfect base pairing of the terminal base pair.** Comparative analysis of PTAP duplication lengths in HIV-1B and HIV-1C. Gag sequences from the LANL database were classified by duplication length, and the prevalence of each length is shown as the percentage of sequences for HIV-1B (N = 6,797, shown in black) and HIV-1C (N = 2,401, shown in grey). The middle panel indicates the presence or absence of a perfect base-pair match between nascent cDNA and the acceptor RNA (color-coded), and the right panel summarizes the number and percentage of sequences with each duplication. Statistical analysis was performed using Pearson’s Chi-Square Test.

Notably, there is no direct evidence on how mismatched bases influence sequence duplication frequencies. Analysis of PTAP motif duplications in p6-Gag provided an opportunity to examine this question, given the substantial variation in duplication length observed in natural sequences. We previously reported that PTAP motif duplication is highly polymorphic, displaying variation in duplication length, sequence divergence between the original and duplicated segments, and subtype-associated patterns (Sharma et al., 2018). For example, HIV-1B preferentially duplicates only three residues (APP) of the six-amino-acid PTAP core motif (EPTAPP), although the biological significance of this partial duplication remains unclear. In contrast, HIV-1C more frequently duplicates six or seven amino acids (EPTAPPA), thereby reproducing the core motif in full. Nevertheless, both subtypes can generate duplications up to 16 amino acids in length (Figure 1). In HIV-1C, additional but smaller peaks are also observed at nine, twelve, and fourteen amino acids (Sharma et al., 2018). Although these data demonstrate subtype-specific preferences in duplication length, the presence of multiple peaks within each subtype suggests that strand-transfer efficiency alone cannot fully account for the observed diversity.

To explore whether sequence features at the strand-switch junction influence duplication outcomes, we compared nucleotide sequences between the original and duplicated motifs. This analysis revealed that peaks in duplication length strongly correlate with perfect base pairing between the 3′ end of the nascent cDNA and the terminal base of the acceptor RNA template (Figure 5C). When three nucleotides of the cDNA form correct base pairs with the acceptor template, an efficient micro-homology domain (MHD) is established, and duplication frequency increases substantially. This pattern is consistently observed at all major duplication peaks in both HIV-1C (six, seven, twelve, and fourteen amino acids) and HIV-1B (three and six amino acids). Thus, complementarity at the 3′ terminus appears to facilitate efficient extension and completion of the duplication event.

In contrast, events requiring mismatch-dependent extension, particularly those lacking an MHD (four, five, eight, nine, ten, eleven, and thirteen amino-acid duplications), occurred at substantially higher frequencies in HIV-1C than in HIV-1B. For example, a nine-amino-acid PTAP motif duplication occurred at a frequency of 1.45% (35/2,401) in HIV-1C, compared with only 0.19% (13/6,797) in HIV-1B (Figure 5C). These observations suggest that HIV-1C RT is more efficient at extending mismatched termini, thereby promoting motif duplication.

To test this hypothesis experimentally, we performed primer extension assays using recombinant RT panels representing HIV-1B and HIV-1C variants to determine whether HIV-1C RT extends mismatched termini more efficiently during reverse transcription. An RNA template corresponding to the PTAP motif region (331 nucleotides; HXB2 coordinates: 1583-1914) was transcribed in vitro. Within this template, we identified a stretch of four bases, ‘AUGC’ (HXB2 coordinates: 1868-1872), where the growing end of primers could align with any one of these four bases (Figure 6A). Four DNA primers were designed with identical sequences except for the terminal nucleotide, such that each primer ended with a different base (A, C, G, or T) at the 3′ end. In each primer set, one primer matched the target template base perfectly, whereas the remaining three represented mismatches. Four such primer sets were generated, each terminating opposite one base of the ‘AUGC’ sequence. Each primer also contained a unique barcode for NGS analysis. In total, the reverse transcription assay employed sixteen primers and three RT enzymes from each subtype panel (bearing G, S, or T at position 359), representing HIV-1B and HIV-1C variants.

**Figure 6.**
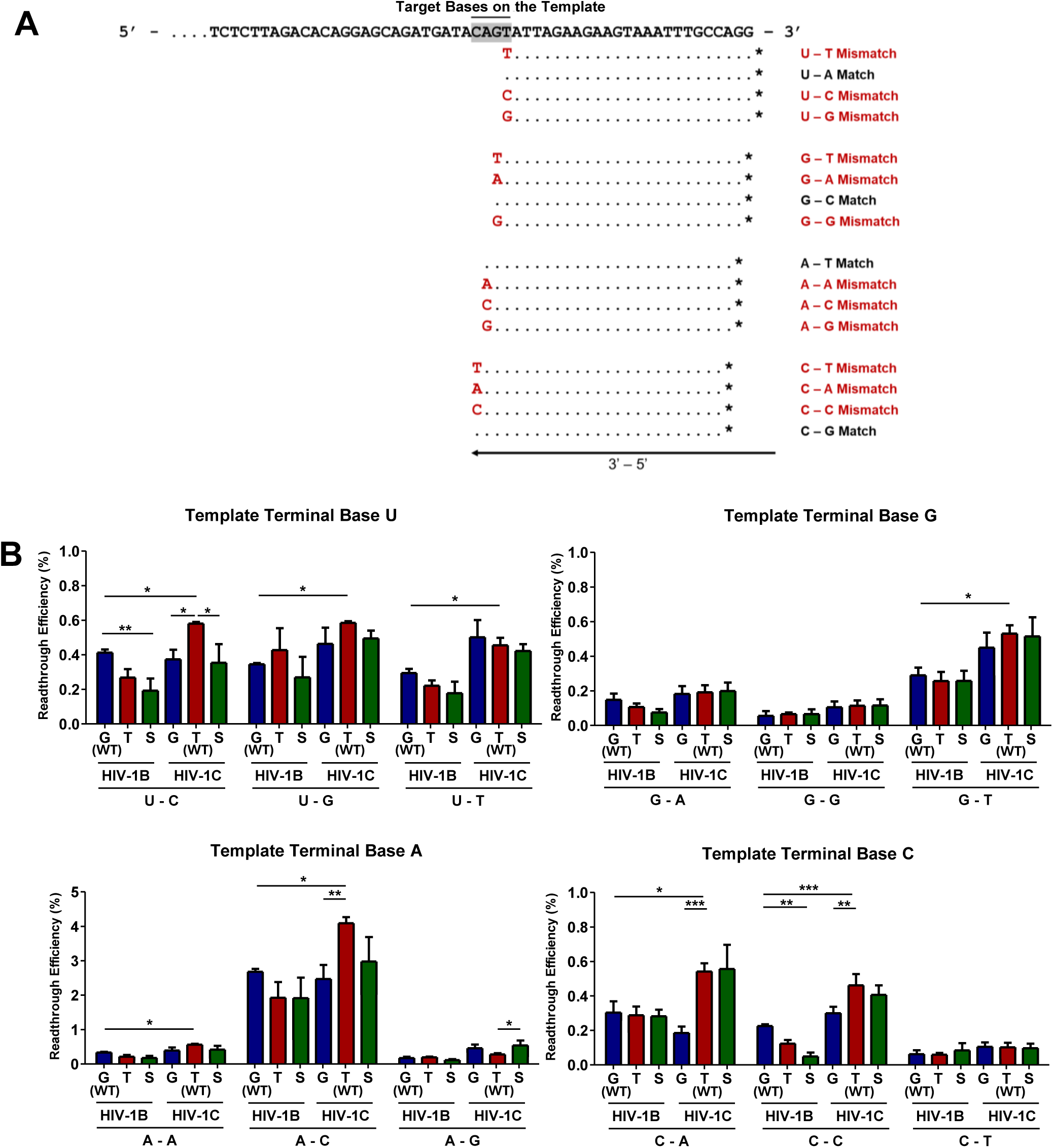
HIV-1C RT can extend mismatched bases more efficiently. (A) Schematic representation of the mismatch extension assay. A 331-nt region on the HIV-1 genome (HXB2: 1583 –1914) was used as template. Four primer sets containing four primers per set each were designed such that three primers in each set carried a distinct 3′ mismatch to represent the 12 possible mismatches; dots indicate complementary bases. The four base pairs where reverse transcription is initiated on the template by the four primer sets is highlighted in grey. * indicates the unique 8 bp barcode for each primer **(B) HIV-1C RT is more efficient at extending mismatched bases.** An RNA template was generated by *in vitro* transcription and annealed to 16 individual barcoded primers representing all base-pair combinations. Reverse transcription was performed with 1 U of the corresponding RT variant, and the efficiency of reverse transcription was estimated by Illumina sequencing. Readthrough efficiency was calculated by normalizing the read counts for the mismatched base pair combinations for each RT with the corresponding matched base read counts and are represented as a percentage. The terminal template base is shown above each panel, with the corresponding primer base indicated below the X-axis. RT variants are color-coded by the amino acid at position 359. Data represent mean ± SD from three independent experiments. Statistical analysis was performed by one-way ANOVA with Tukey–Kramer post hoc test. Statistical significance is shown only for WT comparisons and their corresponding variants and where test results are significant. ***p < 0.001, **p < 0.01, *p < 0.05.

Extension reactions were performed for all primer-template combinations using the RT variants from each panel. The extended products were subjected to second-strand synthesis and prepared for Illumina sequencing to quantify mismatch extension efficiency. Sequencing reads corresponding to each primer-template pair were enumerated, and readthrough efficiency was calculated as the proportion of products in which the mismatched 3′ terminus was successfully extended, normalized to the number of reads obtained for the corresponding matched control primer for each RT variant.

Mismatch extension was observed under all experimental conditions, with readthrough efficiencies ranging from 0.06% to 4.09% (Figure 6B). Consistent with previous reports (Perrino et. al, 1989; Bakhanashvili and Hizi, 1996; Menéndez-Arias, 1998; Kharytonchyk et. al, 2016), purine-purine and pyrimidine-pyrimidine mismatches showed the lowest readthrough efficiencies across RT variants. In contrast, purine-pyrimidine mismatches displayed substantially higher readthrough efficiencies, with an average readthrough of 0.96% across variants. Among these, the A-C mismatch exhibited the highest readthrough potential, followed by U-G, C-A, and G-T.

Overall, HIV-1C RT variants exceeded HIV-1B RTs in extending mismatched bases under most experimental conditions, with many differences reaching statistical significance. The canonical T359 HIV-1C RT showed enhanced extension of eight of the twelve possible mismatch types (A-A, A-C, C-A, C-C, G-T, U-C, U-G, and U-T) compared with the canonical G359 HIV-1B RT. For example, when the terminal base on the template was ‘A’ and the primer ended with a mismatched ‘C’, the readthrough efficiencies of the three HIV-1B RTs (G359, T359, and S359) were 2.69 ± 0.09%, 1.92 ± 0.45%, and 1.91 ± 0.59%, respectively. The corresponding efficiencies for HIV-1C RTs were 2.46 ± 0.41%, 4.09 ± 0.18%, and 2.97 ± 0.71%, respectively. On average, the canonical T359 HIV-1C RT exhibited a 1.6-fold higher mismatch-extension efficiency than the canonical G359 HIV-1B RT. Introducing threonine into HIV-1B RT did not produce statistically significant differences across template-primer combinations. Conversely, substituting glycine (canonical for HIV-1B) for threonine (canonical for HIV-1C) in HIV-1C RT reduced mismatch extension efficiency in only four cases (A-C, C-A, C-C, and U-C). Serine substitutions produced patterns broadly consistent with the respective wild-type residues within each subtype (Figure 6B).

Collectively, these results indicate that HIV-1C RT possesses an enhanced intrinsic capacity to extend mismatched nucleotides relative to HIV-1B RT. This property, together with MHD-mediated annealing, provides a mechanistic explanation for the higher frequency and greater diversity of sequence motif duplications observed in HIV-1C.

## Discussion

### Recombination Frequency Across Subtypes: A Quantitative Similarity

Our previous work showed that HIV-1C generates sequence motif duplications at specific genomic hotspots that confer a replication fitness advantage (Bachu, Yalla, et al., 2012; Sharma et al., 2018). These duplications, particularly in the LTR and p6-Gag regions, arise through nonhomologous recombination mediated by very short microhomology domains, sometimes as small as three bases (Zhang and Temin, 1993; Yin et al., 1997; Panchapakesan and Ranga, 2025). In the present study, we investigated whether subtype-specific properties of RT, particularly those related to recombination, account for the markedly higher frequency of duplications observed at these hotspots, given that RT is the common viral factor involved in both regions.

Recombination plays a central role in RNA virus evolution by promoting genetic diversity and facilitating the removal of deleterious mutations. In retroviruses, recombination rates are particularly high because of their pseudodiploid genomes, the co-packaging of two RNA strands, and the programmed strand-transfer events that occur during reverse transcription. HIV-1 exhibits recombination frequencies at least 10-fold higher than those of other retroviruses, such as Murine Leukaemia Virus and Spleen Necrosis Virus (Onafuwa et al., 2003). Although early studies attributed this high recombination rate primarily to the biochemical properties of RT, subsequent work highlighted the important contributions of RNA packaging and template dynamics (Onafuwa-Nuga and Telesnitsky, 2009).

Whether recombination rates differ among HIV-1 subtypes remains unclear. HIV-1C exhibits a substantially higher frequency of sequence duplications than other subtypes (Bachu, Mukthey, et al., 2012; Bachu, Yalla, et al., 2012; Bhange et al., 2021; Martins et al., 2011; Sharma et al., 2017, 2018), raising the possibility that this subtype undergoes recombination more frequently. However, previous studies (Chin et al., 2005; Galli et al., 2010) and our current data indicate that overall recombination rates are broadly comparable across the two subtypes. Thus, variation in recombination frequency alone cannot explain the duplication bias observed in HIV-1C. Our strand-transfer (Figure 4E) and mismatch-extension (Figure 6B) assays indicate that subtype-specific biochemical properties of HIV-1C RT provide a mechanistic explanation for its elevated duplication frequency.

### HIV-1C RT mediates enhanced strand transfer and mismatch extension events

To our knowledge, no study has directly compared the strand-switching capacity of RTs from different HIV-1 subtypes. Several prior analyses relied on the cell culture-based EGFP complementation assay to measure recombination frequency (Levy et. al, 2004; Rhodes et. al, 2005; Motomura et. al, 2008; Nikolaitchik et. al, 2011). However, this approach lacks the resolution to detect subtle differences in RT strand switching and assesses recombination over only a ∼600 bp region, assuming uniform recombination across the genome. Such an assumption may overlook recombination hotspots and RNA secondary structures. Our study shares this limitation, as additional reporter systems were not evaluated. Nevertheless, because sequence duplication arises from nonhomologous recombination, estimated to occur 10-100 times less frequently than homologous recombination (Zhang & Temin, 1993), the strand-transfer assay provides a relevant surrogate. By quantifying cDNA synthesis on the acceptor strand after a forced transfer, it models conditions that approximate duplication events.

Nonhomologous recombination poses two principal challenges. First, the nascent cDNA-RNA intermediate is inherently unstable due to minimal complementarity between the 3′ end of the DNA and short microhomology regions. We previously proposed that formation of a transient DNA loop constitutes a critical intermediate in duplication (Panchapakesan and Ranga, 2025). Second, terminal mismatches are common, occurring in approximately three out of four events, and have been shown to promote strand transfer (Chin et al., 2007; Palaniappan et al., 1996; Schlub et al., 2014).

Therefore, for duplication to occur, such mismatches must be extended rather than resolved by an additional strand switch, which requires that RT tolerate mismatches and resume polymerisation efficiently. Our data suggest that threonine at position 359 in the connection domain of HIV-1C RT may stabilize the looped intermediate, potentially via an additional hydrogen bond with the nascent DNA. Together with the enhanced mismatch-extension capacity of HIV-1C RT (Figure 6B), this could promote continuous synthesis on the acceptor RNA without further switching, facilitating hotspot-associated duplications (Figure 1C). Notably, the increase in mismatch extension (∼1.6-fold) is modest, consistent with the need to balance tolerance with replication fidelity and avoid excessive error accumulation.

In summary, enhanced strand transfer and improved mismatch extension provide a mechanistic explanation for the elevated frequency of sequence motif duplications in HIV-1C. Further work is needed to define the contribution of additional signature residues to the distinct biochemical properties of HIV-1C RT.

### Signature amino acid residues may modulate the overall performance of the RT

Bioinformatic comparison of RT sequences across HIV-1 subtypes identified several residues unique to HIV-1C (Table 1). We focused on the non-conservative substitution of glycine with threonine at position 359 in the connection domain. Given its proximity to the nascent DNA, threonine could form an additional hydrogen bond with the growing strand. Although the absence of a resolved HIV-1C RT crystal structure limits direct confirmation, two lines of evidence support this hypothesis. First, replacing T359 with glycine, which lacks a hydroxyl group, markedly increased RT activity (Figure 3B), whereas substitution with serine, capable of forming a weaker hydrogen bond, produced only a modest increase. Second, because HIV-1B and HIV-1C RTs share 88–93% sequence identity, structural modelling based on the HIV-1B RT structure reliably predicts hydrogen bond formation at position 359 (Figure 2C).

Consistent with prior reports (Xu et al., 2010), we observed comparable specific activities for wild-type HIV-1B and HIV-1C RTs (Figure 3B). However, substituting glycine for threonine at position 359 increased C-RT activity several-fold, suggesting that the threonine-mediated hydrogen bond constrains polymerase activity. Given the tightly coordinated nature of reverse transcription, excessive RT activity could disrupt replication kinetics. Supporting this interpretation, the threonine-containing variant showed superior p24 production relative to the glycine variant (Figure 3D).

Although our analysis centres on T359, other HIV-1C-specific residues likely contribute to RT function. Notably, C-RT harboring glycine at position 359 displays approximately six-fold higher polymerase activity than B-RT, implying additional cooperative effects. Thus, T359 may act in a compensatory manner to maintain optimal polymerization efficiency. We identified five additional subtype-specific residues, including A36 and T48 near the dNTP-binding pocket in the fingers domain. Because this domain interacts extensively with viral RNA (Patel & Loeb, 2001; Warrilow et al., 2009), variation at these positions may further influence strand transfer and mismatch extension in HIV-1C RT.

### Selection of Duplication Hotspots and Evolutionary Consequences

Darwinian selection strongly shapes HIV-1 evolution by promoting the emergence and persistence of fitter variants. Our findings indicate that although sequence duplications in HIV-1C may arise throughout the genome, they are enriched at discrete hotspots. Because recombination and duplication frequencies are linked, recombination hotspots are expected to overlap with duplication hotspots. However, database analyses identified only four major regions: LTR, p6, env, and nef. Functional studies from our laboratory show that duplications in LTR and p6 confer measurable replication advantages (Bachu, Yalla, et al., 2012; Sharma et al., 2018), consistent with strong positive selection. Duplications in env and nef may similarly enhance fitness, potentially through immune evasion or modulation of pathogenicity, although these remain to be experimentally validated. In contrast, duplications elsewhere are likely deleterious due to frameshifts or disruption of essential motifs and are therefore removed by purifying selection. Thus, while HIV-1C RT may generate duplications broadly, only those that enhance fitness are retained in circulating strains.

This interaction between mutational capacity and selection has important implications for HIV-1 disease management. Subtypes differ in their rates and pathways of antiretroviral resistance acquisition (Garforth et al., 2010; Singh et al., 2014). The global implementation of the WHO ‘Test and Treat’ strategy may intensify selective pressures, favoring compensatory adaptations. PTAP motif duplication, more frequent in HIV-1C, exemplifies such adaptation. Multiple studies link ART initiation to PTAP duplication (Martins et al., 2011, 2015; Peters et al., 2001), and our preliminary data suggest that this duplication may offset fitness costs of resistance mutations. Further, double-PTAP variants appear to have an advantage over single-PTAP strains in mixed infections (Martins et. al, 2011; Sharma et al., 2018). Supporting this, the prevalence of double-PTAP variants in global HIV-1C sequences rose from 17.6% in 1996 to 31% in 2015 (Sharma et al., 2017).

LTR duplications may have even broader clinical consequences. The LTR is highly sensitive to structural changes, as it controls both gene expression and latency. The addition of an extra NF-κB site increases transcriptional activity (Bachu, Yalla, et al., 2012), whereas RBE-III duplication appears to reduce latency reactivation. Our unpublished observations indicate that dual RBE-III variants are refractory to latency-reversing agents, suggesting the potential formation of more stable reservoirs (Bhange and Panchapakesan et. al, 2025).

Collectively, these findings indicate that the elevated duplication frequency in HIV-1C reflects adaptive evolution rather than stochastic replication error. By enhancing fitness, transmission potential, and persistence under therapeutic pressure, this duplication-prone phenotype represents a significant consideration for global HIV control, particularly given the widespread prevalence of HIV-1C.

### Limitations of the Study and Future Directions

The distinctive nature of sequence-motif duplications in HIV-1 presents substantial experimental challenges. These duplications arise from rare, multi-step events during reverse transcription involving nonhomologous strand transfer and template switching. Given reported recombination frequencies of ∼0.03–0.05% per residue across the 9.5 kb genome (Klarmann et al., 2002), and the fact that nonhomologous recombination occurs 100-1,000-fold less frequently than homologous recombination (Zhang & Temin, 1993), the estimated duplication frequency at any specific site is only 0.000015-0.0025%. When combined with the observation that ∼95% of proviruses are defective due to large deletions or lethal mutations (Imamichi et al., 2020; Pollack et al., 2017), direct detection of duplication events in controlled laboratory systems becomes practically challenging.

Consistent with these estimates, prolonged viral culture experiments failed to yield detectable duplication events (data not shown). Serial passaging of the HIV-1C Indie clone for more than four months did not generate variants such as the 4 NF-κB LTR duplication. In vitro attempts are further limited by the predominance of homologous recombination during reverse transcription, which suppresses nonhomologous products. These constraints necessitated reliance on indirect approaches, including comparative sequence analysis, biochemical assays, and computational inference, to investigate the mechanistic basis of increased duplication frequency in HIV-1C.

Accordingly, although our data demonstrate enhanced strand transfer and mismatch extension by HIV-1C RT, we cannot directly link these properties to duplication events in vitro or in vivo. Additional determinants, including nucleocapsid activity, RNA secondary structure, and local sequence context, remain to be systematically examined. Nonetheless, the biochemical differences observed here strongly support a model in which subtype-specific RT features increase the probability of duplication under physiological conditions.

Multiple independent studies, including our own (Bachu, Mukthey, et al., 2012; Bachu, Yalla, et al., 2012; Boullosa et al., 2014; Martins et al., 2011; Sharma et al., 2017, 2018), consistently report an elevated duplication frequency in HIV-1C. Moreover, inter-subtype chimeric RTs, particularly between HIV-1B and HIV-1C, display reduced enzymatic efficiency (Brenner et al., 2006; Iordanskiy et al., 2010; Nagata et al., 2017), underscoring the functional significance of subtype-specific residues. Despite inherent limitations, our study provides mechanistic insight into the molecular determinants underlying this phenotype and identifies candidate amino acid residues that may contribute to duplication propensity. Further investigation is required to define the cooperative effects of additional polymorphisms in shaping this distinctive enzymatic behavior.

## Materials and Methods

### Sequence and Structural Analysis

HIV-1 sequences were retrieved from the Los Alamos National Laboratory HIV Sequence Database and aligned using Clustal Omega with default parameters. Multiple sequence alignments were examined and curated using BioEdit version 7.2.5. Structural visualization and modelling of the RT were performed using PyMOL version 2.4.1. A high-resolution crystal structure of HIV-1B RT (PDB ID: 5J2M) was used as the template. HIV-1C RT–specific signature amino acid residues were introduced into this structure using the mutagenesis tool in PyMOL, followed by local energy minimization to optimize side-chain conformations. The bond length between the T359 residue and the DNA phosphate (PO₄) group was measured using the built-in distance calculation tools in the software.

### Construction of Plasmids

In all experiments, we used the NL4-3 (GenBank accession number: AF324493.2) and Indie-C1 (AB023804.1) molecular clones as representatives of HIV-1B and -1C, respectively. The NL4-3 and NL4-3-ΔEnv-EGFP molecular clones were obtained through the NIH-AIDS Reagent Program of the Division of AIDS, NIAID, NIH (Cat. Nos: ARP-2852 and ARP-11100, contributed by Dr. M. Martin, and Dr. Haili Zhang, Dr. Yan Zhou, and Dr. Robert Siliciano, respectively). The Indie-C1 molecular clone was a kind gift from Dr. Masashi Tatsumi (International Medical Center of Japan, Tokyo). The Indie-ΔEnv-EGFP reporter vector was generated in our laboratory. Prof. Vinayaka Prasad (Albert Einstein College of Medicine, New York) kindly provided the RT expression vectors p6HRT and p6HRT-p51, originally developed by Stuart LeGrice and Fiona Gruninger-Leitch (1990). All plasmid constructs were cloned in *E. coli* XL-10 Gold® cells, grown at 30°C overnight. All primers were synthesized by Sigma-Aldrich, Bengaluru, India, and enzymes for cloning and PCR were procured from New England Biolabs (USA). Additional reagents used in specific assays are indicated where relevant.

Panels of full-length infectious molecular clones were generated for both NL4-3 and Indie-C1 backbones, differing by a single amino acid at position 359 of RT (glycine, threonine, serine). Site-directed mutagenesis was performed using overlap-extension PCR with Phusion™ DNA polymerase and primer sets listed in Supplementary File 1. Mutant fragments were assembled using outer primer pairs, digested with AgeI/EcoRI (for NL4-3) or PflMI (for Indie-C1), and substituted into the parental backbones via restriction enzyme–mediated fragment exchange. Recombinant clones were identified by restriction digestion and confirmed by Sanger sequencing. The resulting constructs were designated according to the amino acid at position 359 (e.g., Indie–359G).

The RT expression vectors p6HRT (p66 subunit) and p6HRT-p51 (p51 subunit) contain an N-terminal 6×His tag under the control of the lac promoter. The corresponding NL4-3 or Indie RT variants were cloned into the p6HRT and p6HRT-p51 backbones using BamHI/SalI and BamHI/HindIII restriction sites, respectively. A unique restriction site was incorporated into the reverse primer beyond the stop codon to serve as a surrogate identifier for the amino acid at position 359. For the Indie constructs, an additional SacI site was introduced for differentiation from the NL4-3 panel.

Reporter constructs were derived from NL4-3 ΔEnv EGFP and Indie ΔEnv EGFP backbones. The RT region was modified by swapping restriction fragments (AgeI/EcoRI for NL4-3 and PflMI for Indie) from the corresponding full-length RT mutant clones. Each of the eight intermediate RT variant reporters was then used to generate two EGFP frameshift mutants, carrying insertions at amino acid positions 4 or 204 of the GFP open reading frame. Mutations were introduced by direct or overlap PCR and cloned directionally using EcoRI/NheI (for NL4-3) or SphI/StuI (for Indie) restriction sites. Recombinant clones were verified by restriction digestion and confirmed by Sanger sequencing.

### Expression and Purification of Recombinant RTs

Recombinant HIV-1 RT expression constructs (p6HRT and p6HRT-p51) were transformed separately into E. coli M15 competent cells (Stuart & Friona, 1990). Protein expression was induced with 1 mM IPTG at an OD₆₀₀ of 0.4 and continued for 5 h at 37°C with shaking. Cells were harvested, resuspended in 50 mM sodium phosphate buffer (pH 8.0) containing 150 mM NaCl and 1 mM PMSF, and lysed by sonication (VCX-130, Sonics and Materials Inc., USA). The clarified lysate (12,000 × g, 10 min) was incubated overnight at 4°C with Ni–NTA resin (0.5 ml resin/200 ml lysate; G-Biosciences, USA).

The lysate was then washed with 50 mM sodium phosphate (pH 8.0), 20 mM imidazole, 150 mM NaCl, 1 mM PMSF, and the bound RT was eluted with 250 mM imidazole in the same buffer. Eluted fractions were pooled and dialyzed overnight against 50 mM Tris-Cl (pH 7.0), 25 mM NaCl, 1 mM EDTA, 20% glycerol. Further purification was achieved using DEAE-Sepharose ion-exchange chromatography, collecting the flow-through fraction containing RT. Final enzyme preparations were dialyzed into 50 mM Tris-Cl (pH 8.0), 150 mM NaCl, 1 mM DTT, 50% glycerol (Stahlhut & Olsen, 1996), concentrated using Amicon® concentrators (Merck-Millipore, USA), and stored in aliquots at −20°C until use.

### Determination of RT activity

RT activity was assayed using a poly(U)/oligo(dA)₍₂₀₎ template–primer system as described previously (Stuart, Le Grice, & Cameron, 1995), with minor modifications. Briefly, 4 µg polyuridylic acid (Sigma-Aldrich, USA) was annealed with 1 µg (∼80 pmol) oligo(dA)₍₂₀₎ primer by heating at 95°C followed by rapid cooling on ice. The template–primer complex was incubated in a 20 µl reaction containing 25 mM Tris-Cl (pH 8.0), 3 mM MgCl₂, 100 mM KCl, and 250 µM dATP (GeNei Labs, India), supplemented with 1 µCi [α-³²P]-dATP (Board of Radiation & Isotope Technologies, India).

Reactions were initiated with varying enzyme concentrations (0.1–1 µl) and incubated at 37°C for 10 min, then terminated with 0.5 M EDTA. Aliquots were spotted on Hybond-N⁺ nylon membranes (Amersham Lifesciences, USA), washed to remove unincorporated nucleotides, ethanol-rinsed, and air-dried. Radioactivity incorporated into cDNA was quantified by liquid scintillation counting (MicroBeta², PerkinElmer, USA) using Ultima Gold™ XR scintillation fluid.

Specific activity was calculated as the amount of enzyme incorporating 1 pmol [α-³²P]-dATP in 10 min at 37°C per µg of protein. For rate determination, reactions were performed as above with 1 µl enzyme, and nucleotide incorporation was measured at 5-minute intervals.

### Cell Culture, Virus Production, and Titration

T-cell lines CEM-CCR5, Jurkat, Jurkat-CCR5, and epithelial cell lines HEK 293T and TZM-bl, were maintained in RPMI 1640 or DMEM, respectively (Sigma-Aldrich, USA) supplemented with 10% FBS (Life Technologies, India), 2 mM Glutamine, and 100 µg/ml each of penicillin G and streptomycin (Sigma-Aldrich, USA). All cell cultures were incubated at 37°C in 5% CO₂. Peripheral Blood Mononuclear Cells (PBMCs) were isolated from fresh donor blood by density gradient centrifugation. CD8⁺ cells were depleted using the RosetteSep™ Human CD8 Depletion Cocktail (Stemcell Technologies, Canada). PBMCs were activated for 72 h in RPMI 1640 containing 20 U/ml IL-2, 5 µg/ml Phytohaemagglutinin-P (PHA-P), and the supplements described above. Prior to infection, cells were maintained without PHA-P. Infectious HIV-1 viral stocks were generated by transient transfection of HEK 293T cells. Approximately 3 × 10⁶ cells were seeded in 100 mm dishes and transfected with 20 µg of molecular clone DNA and 30 ng of pCMV-TdTomato (transfection control) using the calcium phosphate method (Kingston et al., 2003). For recombination assays, pseudotyped viral variants were produced in 6-well plates by transfecting 0.3 × 10⁶ HEK 293T cells with a mixture containing 3 µg of reporter vector, 1 µg of pCMV-VSV-G, and 10 ng of pCMV-TdTomato. Viruses copackaging two RNA genomes, each harboring complementary debilitating mutations in the GFP ORF, were produced by cotransfecting equal amounts (1.5 µg each) of the corresponding plasmids. Following transfection, the culture medium was replaced after 6 h, and supernatants were harvested at 48 h post-transfection, filtered through 0.22 µm syringe filters, and stored in aliquots at −80°C. Viral yield was quantified by p24 ELISA (4th Generation HIV-1 p24 Antigen ELISA Kit, J. Mitra & Co. Pvt. Ltd., India). Functional infectivity was assessed by infecting TZM-bl cells with serial dilutions of the viral stock. Briefly, 1 × 10⁴ cells were seeded per well in 96-well plates and infected in the presence of 10 µg/ml DEAE-dextran. After 8 h, the inoculum was replaced with complete DMEM, and cells were incubated for 48 h at 37°C and 5% CO₂. Infection efficiency was quantified by measuring firefly luciferase activity in cell lysates using the Luciferase Assay System (Promega, USA) on a Varioskan Lux multimode reader (Thermo Fisher Scientific, USA).

### Recombination Assay

CEM-CCR5 or Jurkat cells (0.3 × 10⁶) were infected with 10 ng p24 equivalent of either NL4-3 or Indie ΔEnv EGFP variant viruses in 2 ml RPMI medium containing 10 µg/ml DEAE-dextran in 6-well plates. The culture medium was replaced with fresh RPMI 6 h post-infection. Twenty-four hours before analysis, cells were activated with a cocktail containing 10 ng/ml TNF-α (Miltenyi Biotec, USA), 5 ng/ml PMA, and 5 mM HMBA (Sigma-Aldrich, USA). Cells were harvested by centrifugation at 500 × g, washed twice with PBS, and resuspended in 250 µl PBS with 2% FBS. Flow cytometric analysis was performed using a BD FACS ARIA III (Becton, Dickinson and Company, USA) to quantify GFP expression as a measure of recombination frequency.

### Strand-Transfer Assay

Two regions of the Indie genome (HXB2 coordinates 256–462 and 473–603) were amplified using forward primers carrying a T7 promoter (Supplementary File 1) to enable in vitro transcription. The second fragment contained a 20 bp sequence at its 3′ end homologous to the 5′ end of the first fragment, allowing potential strand transfer during reverse transcription. RNAs were synthesized and purified, mixed at an equimolar ratio, and reverse transcribed using 1 Unit of RT and the antisense primer N4864, which anneals to RNA-1, under the same conditions. Two µl of the cDNA product were diluted 1,000-fold and used as templates for two qPCR reactions: one with primers N4863/N4864 to detect the RNA-1 template, and another with N4865/N4864 to detect strand-transfer products.

### Mismatch Extension Assay

A 331-nt fragment of the HIV-1C genome (HXB2 coordinates: 1583–1914) was PCR-amplified using primers containing the T7 promoter sequence to enable in vitro transcription. The resulting RNA served as the template, with a defined four-base stretch (ATGC) used as the initiation site for polymerization (Figure 6A). Sixteen reverse primers, grouped by an initiation base, were designed so that each group initiated synthesis at one of the four template bases. Within each group, primers were identical except for the 3′ terminal nucleotide, yielding all sixteen possible base pair combinations at the primer–template junction, four correct (A–T, T–A, G–C, C–G) and twelve mismatched pairs. Three independent sets of the sixteen primers were synthesized, each carrying a unique 8 bp barcode at the 5′ end to enable multiplexed next-generation sequencing (NGS). Primer sequences are listed in Supplementary File 1. Annealing was performed by mixing 2 µg of template with 100 pmol of the appropriate primer, and the extension reactions were carried out using the six recombinant RT variants under identical conditions. Reaction products were treated with RNase H, purified, and subjected to second-strand synthesis using Taq DNA polymerase. Products were then purified, and equal reaction volumes were pooled for end repair and library preparation, followed by paired-end Illumina sequencing on a NextSeq 2000 platform. Sequencing data were analyzed using a custom Python pipeline.

## Acknowledgements

We thank Prof. Vinayaka Prasad for the critical reading of the manuscript.

---

**Supplementary Table 1:**
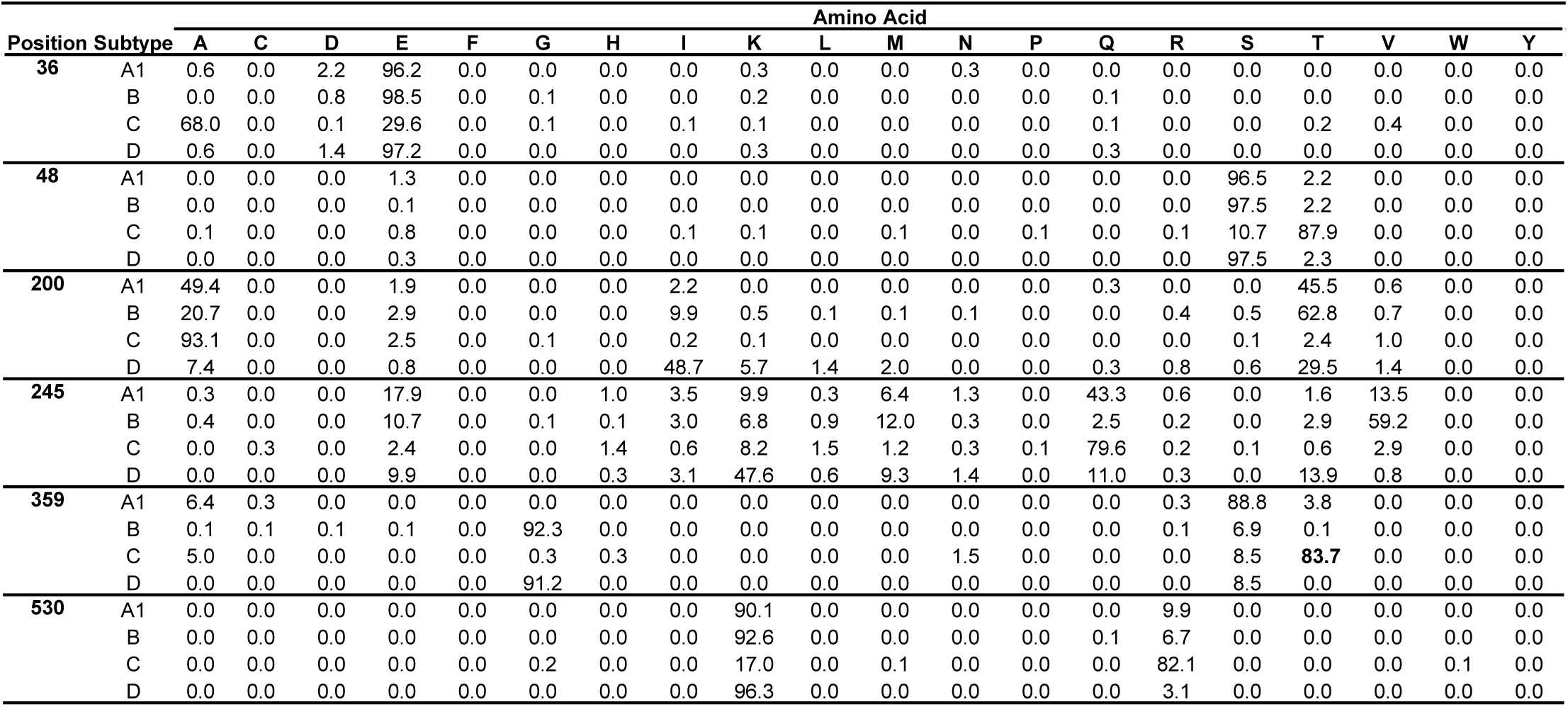
The percentage prevalence of amino acids in various subtypes at the six locations shown in Figure 2A. The single letter amino acid code is used.

|  |  | Amino Acid |  |  |  |  |  |  |  |  |  |  |  |  |  |  |  |  |  |  |  |
| --- | --- | --- | --- | --- | --- | --- | --- | --- | --- | --- | --- | --- | --- | --- | --- | --- | --- | --- | --- | --- | --- |
| Position | Subtype | A | C | D | E | F | G | H | I | K | L | M | N | P | Q | R | S | T | V | W | Y |
| 36 | A1 | 0.6 | 0.0 | 2.2 | 96.2 | 0.0 | 0.0 | 0.0 | 0.0 | 0.3 | 0.0 | 0.0 | 0.3 | 0.0 | 0.0 | 0.0 | 0.0 | 0.0 | 0.0 | 0.0 | 0.0 |
|  | B | 0.0 | 0.0 | 0.8 | 98.5 | 0.0 | 0.1 | 0.0 | 0.0 | 0.2 | 0.0 | 0.0 | 0.0 | 0.0 | 0.1 | 0.0 | 0.0 | 0.0 | 0.0 | 0.0 | 0.0 |
|  | C | 68.0 | 0.0 | 0.1 | 29.6 | 0.0 | 0.1 | 0.0 | 0.1 | 0.1 | 0.0 | 0.0 | 0.0 | 0.0 | 0.1 | 0.0 | 0.0 | 0.2 | 0.4 | 0.0 | 0.0 |
|  | D | 0.6 | 0.0 | 1.4 | 97.2 | 0.0 | 0.0 | 0.0 | 0.0 | 0.3 | 0.0 | 0.0 | 0.0 | 0.0 | 0.3 | 0.0 | 0.0 | 0.0 | 0.0 | 0.0 | 0.0 |
| 48 | A1 | 0.0 | 0.0 | 0.0 | 1.3 | 0.0 | 0.0 | 0.0 | 0.0 | 0.0 | 0.0 | 0.0 | 0.0 | 0.0 | 0.0 | 0.0 | 96.5 | 2.2 | 0.0 | 0.0 | 0.0 |
|  | B | 0.0 | 0.0 | 0.0 | 0.1 | 0.0 | 0.0 | 0.0 | 0.0 | 0.0 | 0.0 | 0.0 | 0.0 | 0.0 | 0.0 | 0.0 | 97.5 | 2.2 | 0.0 | 0.0 | 0.0 |
|  | C | 0.1 | 0.0 | 0.0 | 0.8 | 0.0 | 0.0 | 0.0 | 0.1 | 0.1 | 0.0 | 0.1 | 0.0 | 0.1 | 0.0 | 0.1 | 10.7 | 87.9 | 0.0 | 0.0 | 0.0 |
|  | D | 0.0 | 0.0 | 0.0 | 0.3 | 0.0 | 0.0 | 0.0 | 0.0 | 0.0 | 0.0 | 0.0 | 0.0 | 0.0 | 0.0 | 0.0 | 97.5 | 2.3 | 0.0 | 0.0 | 0.0 |
| 200 | A1 | 49.4 | 0.0 | 0.0 | 1.9 | 0.0 | 0.0 | 0.0 | 2.2 | 0.0 | 0.0 | 0.0 | 0.0 | 0.0 | 0.3 | 0.0 | 0.0 | 45.5 | 0.6 | 0.0 | 0.0 |
|  | B | 20.7 | 0.0 | 0.0 | 2.9 | 0.0 | 0.0 | 0.0 | 9.9 | 0.5 | 0.1 | 0.1 | 0.1 | 0.0 | 0.0 | 0.4 | 0.5 | 62.8 | 0.7 | 0.0 | 0.0 |
|  | C | 93.1 | 0.0 | 0.0 | 2.5 | 0.0 | 0.1 | 0.0 | 0.2 | 0.1 | 0.0 | 0.0 | 0.0 | 0.0 | 0.0 | 0.0 | 0.1 | 2.4 | 1.0 | 0.0 | 0.0 |
|  | D | 7.4 | 0.0 | 0.0 | 0.8 | 0.0 | 0.0 | 0.0 | 48.7 | 5.7 | 1.4 | 2.0 | 0.0 | 0.0 | 0.3 | 0.8 | 0.6 | 29.5 | 1.4 | 0.0 | 0.0 |
| 245 | A1 | 0.3 | 0.0 | 0.0 | 17.9 | 0.0 | 0.0 | 1.0 | 3.5 | 9.9 | 0.3 | 6.4 | 1.3 | 0.0 | 43.3 | 0.6 | 0.0 | 1.6 | 13.5 | 0.0 | 0.0 |
|  | B | 0.4 | 0.0 | 0.0 | 10.7 | 0.0 | 0.1 | 0.1 | 3.0 | 6.8 | 0.9 | 12.0 | 0.3 | 0.0 | 2.5 | 0.2 | 0.0 | 2.9 | 59.2 | 0.0 | 0.0 |
|  | C | 0.0 | 0.3 | 0.0 | 2.4 | 0.0 | 0.0 | 1.4 | 0.6 | 8.2 | 1.5 | 1.2 | 0.3 | 0.1 | 79.6 | 0.2 | 0.1 | 0.6 | 2.9 | 0.0 | 0.0 |
|  | D | 0.0 | 0.0 | 0.0 | 9.9 | 0.0 | 0.0 | 0.3 | 3.1 | 47.6 | 0.6 | 9.3 | 1.4 | 0.0 | 11.0 | 0.3 | 0.0 | 13.9 | 0.8 | 0.0 | 0.0 |
| 359 | A1 | 6.4 | 0.3 | 0.0 | 0.0 | 0.0 | 0.0 | 0.0 | 0.0 | 0.0 | 0.0 | 0.0 | 0.0 | 0.0 | 0.0 | 0.3 | 88.8 | 3.8 | 0.0 | 0.0 | 0.0 |
|  | B | 0.1 | 0.1 | 0.1 | 0.1 | 0.0 | 92.3 | 0.0 | 0.0 | 0.0 | 0.0 | 0.0 | 0.0 | 0.0 | 0.0 | 0.1 | 6.9 | 0.1 | 0.0 | 0.0 | 0.0 |
|  | C | 5.0 | 0.0 | 0.0 | 0.0 | 0.0 | 0.3 | 0.3 | 0.0 | 0.0 | 0.0 | 0.0 | 1.5 | 0.0 | 0.0 | 0.0 | 8.5 | 83.7 | 0.0 | 0.0 | 0.0 |
|  | D | 0.0 | 0.0 | 0.0 | 0.0 | 0.0 | 91.2 | 0.0 | 0.0 | 0.0 | 0.0 | 0.0 | 0.0 | 0.0 | 0.0 | 0.0 | 8.5 | 0.0 | 0.0 | 0.0 | 0.0 |
| 530 | A1 | 0.0 | 0.0 | 0.0 | 0.0 | 0.0 | 0.0 | 0.0 | 0.0 | 90.1 | 0.0 | 0.0 | 0.0 | 0.0 | 0.0 | 9.9 | 0.0 | 0.0 | 0.0 | 0.0 | 0.0 |
|  | B | 0.0 | 0.0 | 0.0 | 0.0 | 0.0 | 0.0 | 0.0 | 0.0 | 92.6 | 0.0 | 0.0 | 0.0 | 0.0 | 0.1 | 6.7 | 0.0 | 0.0 | 0.0 | 0.0 | 0.0 |
|  | C | 0.0 | 0.0 | 0.0 | 0.0 | 0.0 | 0.2 | 0.0 | 0.0 | 17.0 | 0.0 | 0.1 | 0.0 | 0.0 | 0.0 | 82.1 | 0.0 | 0.0 | 0.0 | 0.1 | 0.0 |
|  | D | 0.0 | 0.0 | 0.0 | 0.0 | 0.0 | 0.0 | 0.0 | 0.0 | 96.3 | 0.0 | 0.0 | 0.0 | 0.0 | 0.0 | 3.1 | 0.0 | 0.0 | 0.0 | 0.0 | 0.0 |

**Supplementary Table 2:**
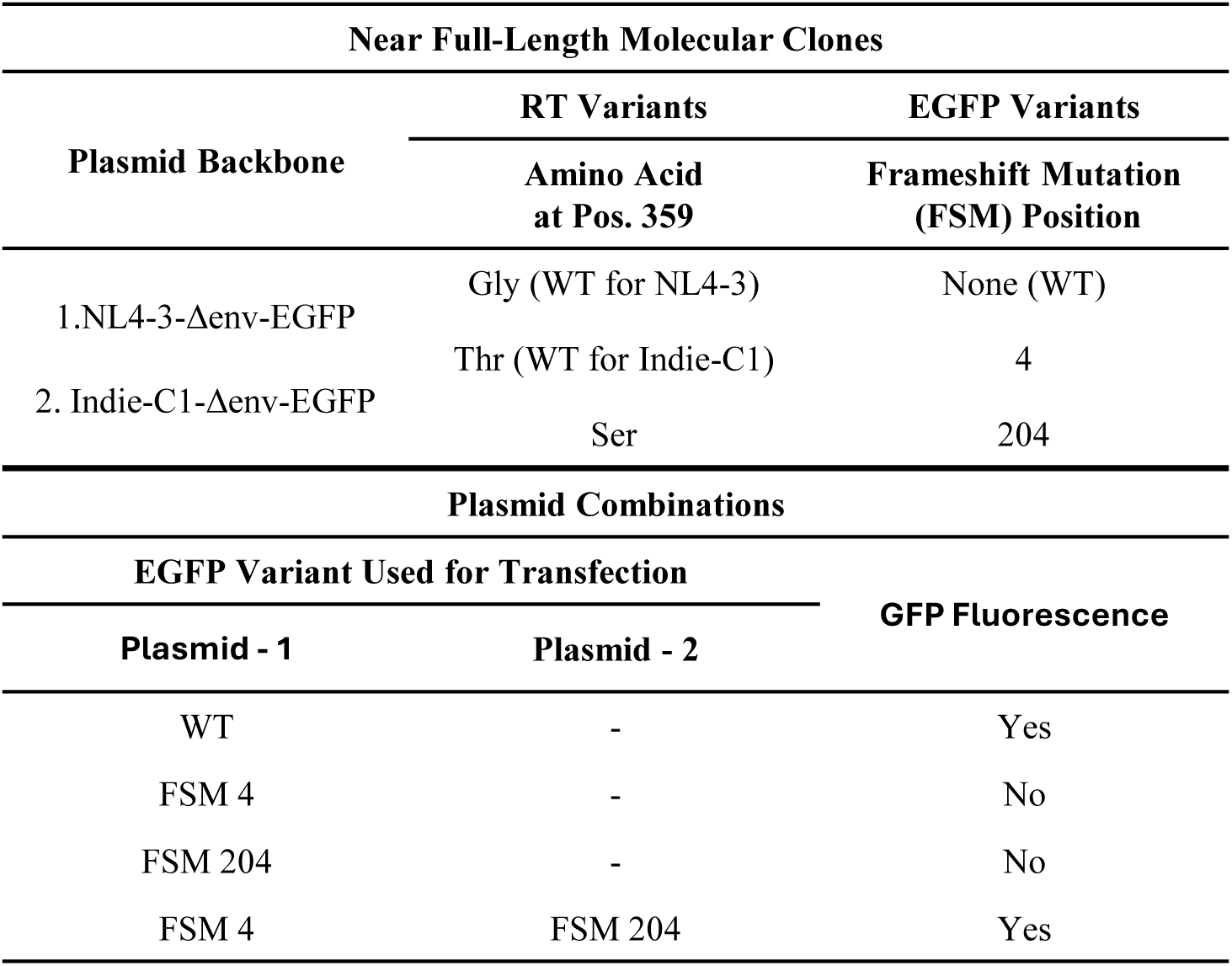
The composition of the viral variants shown in Figure 4A and the plasmid combinations that were used for virus production.

